# Galectin-3 C-epitope, ANP, PPIA, and albumin function as drug-responsive panel biomarkers and therapeutic targets in aging and pressure overload-induced cardiac hypertrophy

**DOI:** 10.1101/2025.05.31.657024

**Authors:** Puja Laxmanrao Shinde, Vikas Kumar, Siddhartha Singh, K.C. Sivakumar, Rashmi Mishra

## Abstract

Biological aging (BA) and pressure overload-induced cardiac hypertrophy (PO-CH) are marked by myocardial thickening, fibrosis, and functional decline, culminating in an increased risk of heart failure. Effective clinical management requires biomarkers that not only track disease progression but also monitor therapeutic response. Using Amalaki Rasayana (AR), a cardioprotective nutraceutical-based medicine, in rat models of BA and PO-CH, we identified a panel of serum biomarkers whose treatment-induced decline was associated with disease regression. Among them, galectin-3 emerged in both full-length and oligomeric C-epitope cleaved forms, with the latter closely linked to pathological surface remodeling. Treatment with AR and its bioactive component gallic acid (GA) suppressed extracellular galectin-3 C-epitope oligomer accumulation by inhibiting full-length galectin-3 secretion through phosphorylation-dependent mechanisms and promotion of intracellular retention. Concurrently, serum levels of atrial natriuretic peptide (ANP), cyclophilin A (PPIA), and albumin (ALB) were consistently modulated by therapy. Validation in sera from elderly individuals and cardiac hypertrophy patients responding to conventional treatments supported the translational relevance of this biomarker panel. These findings establish a novel, mechanistically grounded biomarker set—galectin-3 C-epitope, ANP, PPIA, and ALB—for monitoring therapeutic response in cardiac hypertrophy, and position AR as a promising medicinal intervention against age- and pressure-induced cardiac pathology.

**GRAPHICAL ABSTRACT:** 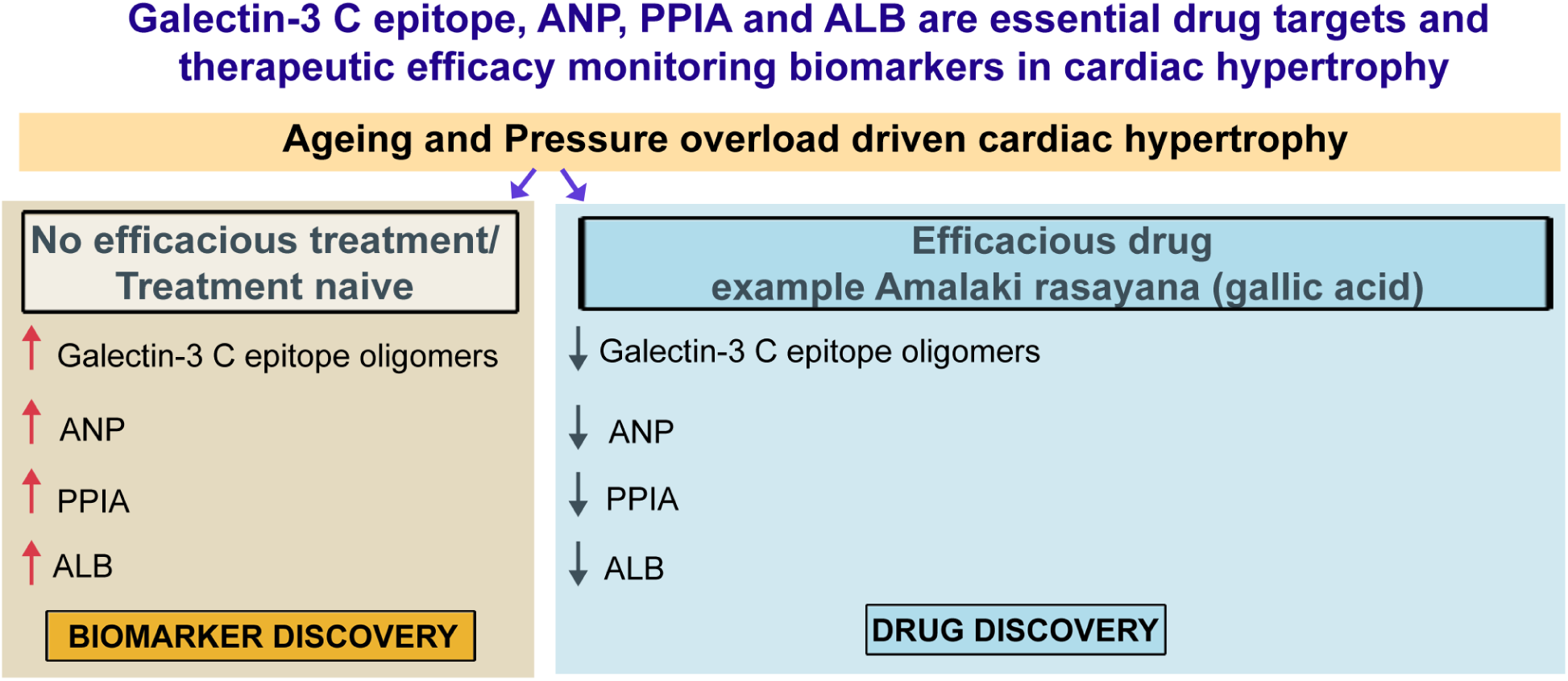

**HIGHLIGHTS:**

- Galectin-3 C-epitope, ANP, PPIA, and ALB are elevated in aging and pressure overload driven left ventricular hypertrophy.
- Amalaki rasayana (AR) and Gallic Acid (GA) reduce these markers in cardiac tissue and circulation.
- AR and GA inhibit galectin-3 secretion and oligomeric binding to cardiomyocyte surfaces.
- AR and GA regulation predominantly occurs via phosphorylation-dependent cytoplasmic retention of galectin-3.
- These markers serve as potential therapeutic targets and diagnostic biomarkers.

## INTRODUCTION

Hypertrophic cardiomyopathy (HCM), primarily driven by pressure overload in the left ventricle, and its advanced stage, dilated cardiomyopathy (D-HCM), represent major contributors to heart failure characterized by left ventricular systolic dysfunction and chamber dilatation [1, 2]. Initially, cardiomyocytes respond to elevated mechanical stress— arising from increased intraventricular pressure and a stiffened extracellular matrix—by hypertrophic growth, increasing in mass to maintain cardiac output [2]. However, chronic pressure overload eventually precipitates cardiomyocyte thinning, stretching, cellular damage, and death, culminating in D-HCM and progressive cardiac dysfunction [3]. Globally, the rising incidence of hospitalizations, readmissions, and mortality related to HCM and D-HCM highlights the urgent need for sensitive diagnostic and prognostic biomarkers that reflect disease progression and recovery [4, 5, 6].

Effective biomarkers should dynamically correlate with disease severity, increasing during exacerbation and declining with recovery, thereby serving not only as indicators but also as therapeutic targets. Importantly, biomarker-guided drug therapy enables clinicians to tailor treatment regimens—optimizing drug choice, dosage, and duration—and ultimately improving patient outcomes [5]. Such ideal biomarkers must possess high sensitivity, specificity, stability, cost-effectiveness, and minimal invasiveness, while maintaining reliability across diverse populations and sexes [6].

Myocardial fibrosis is a key pathological driver of hypertrophic and dilated cardiac failure. Fibrotic remodeling increases left ventricular stiffness, promotes ventricular dyssynchrony and tissue heterogeneity, and elevates cardiac afterload, thereby escalating the risk of heart failure and sudden cardiac death [5, 6]. Consequently, biomarkers and drug targets emerging from the cardiac fibrotic proteome hold considerable promise. Among these, galectin-3 (gal- 3)—a beta-galactoside-binding lectin—has garnered attention due to its strong association with myocardial fibrosis, left ventricular dysfunction, and adverse cardiac outcomes [5, 6, 7]. Elevated circulating galectin-3 levels correlate with enhanced fibrosis, as gal-3 actively promotes fibroblast activation and scar formation in injured myocardium [8, 9]. The FDA’s approval of galectin-3 assays for prognostic use in heart failure further validates its clinical utility in predicting mortality, hospitalization, and therapeutic response, with consistent performance across sexes and geographic regions [10, 11, 12].

Galectin-3 comprises two functional domains: an N-terminal domain (∼110-115 amino acids, Gal-3N) and a C-terminal carbohydrate recognition domain (∼130-150 amino acids, Gal-3C) [13]. This protein exists both as a full-length molecule (26–35 kDa) and in cleaved forms— Gal-3N (∼15.5 kDa) and Gal-3C (∼10.5 kDa)—likely generated extracellularly through proteolytic cleavage by collagenases and matrix metalloproteinases [13].

The C-terminal domain (also called C-epitope) mediates glycan binding and oligomerization, facilitating significant signaling modulation and bioactivation [14, 15]. In cardiac tissue, galectin-3 C-terminal oligomers predominantly interact with glycosylated surface receptors on fibroblasts, macrophages, endothelial cells, and cardiomyocytes, linking galectin-3’s molecular functions to crucial cellular processes in cardiac remodeling and disease [16]. We hypothesized that aberrant or elevated circulating levels of galectin-3 C-epitope oligomers could serve as superior biomarkers to inform therapeutic strategies in HCM and D-HCM.

Our objective was to demonstrate that therapeutic interventions targeting cardiac hypertrophy should focus on reducing the pool of galectin-3 C-epitope oligomers. By quantifying circulating gal-3C oligomers and monitoring their modulation during treatment, we aimed to correlate these levels with the progression or regression of hypertrophic cardiac damage. To test this, an in vivo drug model capable of alleviating left ventricular hypertrophy was essential to differentiate disease from recovery phases.

Amalaki rasayana (AR), a traditional Indian medicine derived from *Phyllanthus <u>emblica</u>* fruit, has gained recognition for its efficacy against biological and pathological cardiac hypertrophy [17, 18, 19]. Noted for its potent antioxidant, anti-fibrotic, and anti-aging properties, recent multi-omics studies have reinforced AR’s role as a promising anti-cardiac hypertrophy agent [18, 19, 20]. AR is rich in gallic acid (GA) and its metabolite ellagic acid, both shown to confer cardioprotection via mechanisms including mitochondrial health enhancement [18, 21, 22]. Untargeted serum proteomics in AR-treated rat models have identified significant GA circulation, suggesting direct targeting of cardiac hypertrophy rather than mere metabolic by-products like ellagic acid or urolithins [19]. Moreover, Bergenin—a natural GA derivative—has demonstrated in vitro inhibition of galectin-3 in cancer cell cultures, highlighting GA’s therapeutic potential in regulating galectin-3 homeostasis [23, 24].

Using AR-based animal models of biological aging and pressure overload-induced pathological cardiac hypertrophy, we sought to elucidate the significance of galectin-3 C-epitope oligomers. We aimed to establish their superiority over full-length galectin-3 in evaluating drug efficacy against aging and pressure overload-driven left ventricular hypertrophy. Our findings reveal a promising biomarker signature for monitoring cardiac hypertrophy treatment, detailed in the results section. Given the global burden of hypertrophic cardiac diseases, this research addresses a critical unmet need with the potential to transform therapeutic approaches for millions worldwide.

## RESULTS

### Amalaki rasayana (AR) significantly reduces extracellular and circulating galectin-3 C-epitope oligomers in a rat model of left ventricular cardiac hypertrophy (CH)

To evaluate the therapeutic potential of galectin-3 C-epitope as both a biomarker and drug target for cardiac hypertrophy (CH), we developed an in vivo Wistar rat model of CH induced by biological aging (BA) and pressure overload (PO). Recognizing the increasing interest in complementary and integrative medicine for cardiovascular health, we selected Amalaki rasayana (AR), an Ayurvedic nutraceutical with established antioxidant, anti-inflammatory, and cardioprotective properties, as the intervention agent [17, 18].

Rats were administered AR orally at a dosage of 500 mg/kg body weight, following previously established protocols [18]. Over the 21-month experimental period, we assessed a comprehensive range of cardiac function parameters to determine AR’s efficacy (see experimental design, **Figure 1A).** Compared with untreated BA and PO-CH controls, AR-treated cohorts exhibited significant improvement in echocardiographic indices, including left ventricular fractional shortening (LVFS) and ejection fraction (LVEF) **(Figure 1B–D)**.

**Figure 1.**
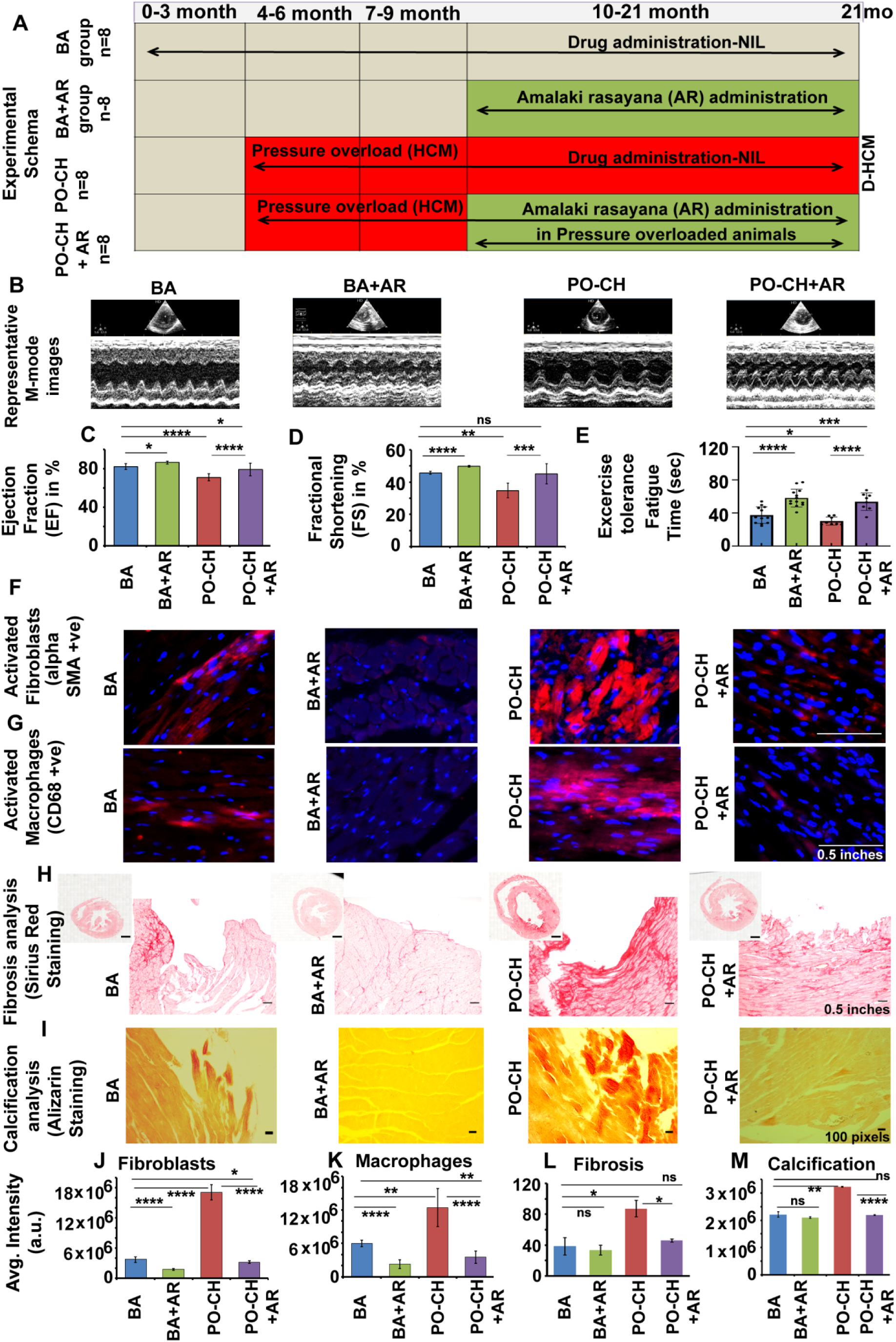
Development and validation of an Amalaki rasayana (AR)-based therapeutic model for left ventricular cardiac hypertrophy (CH). **(A)** Experimental design: Age-matched rat groups included BA (21-month-old biologically aged rats with mild age-related CH), BA+AR (BA rats treated with AR from 9 to 21 months, totalling 12 months), PO-CH (rats subjected to trans-aortic constriction at 3^rd^ month, developing pressure overload-induced CH by 9^th^ month and ventricular dilation by 21 months), and PO-CH+AR (PO-CH rats treated with AR for ≥12 months post-surgery, from 9^th^ to 21^st^ month). **(B–D)** M-mode echocardiography assessing left ventricular ejection fraction (LVEF), fractional shortening (LVFS), and ventricular dimensions showed significant improvement in AR-treated groups compared to untreated BA and PO-CH cohorts. **(E)** Exercise tolerance tests measuring fatigue time indicated enhanced cardiac output in AR-treated rats versus untreated groups. **(F, G, J, K)** Immunohistochemistry for CD68+ve enlarged macrophages and αSMA over-expressing activated fibroblasts revealed significant reductions in AR-treated animals relative to controls. **(H, L)** Sirius red staining demonstrated decreased myocardial fibrosis in AR-treated groups. **(I, M)** Alizarin S staining indicated reduced myocardial calcification with AR treatment. Data represent mean ± SD; significance: *p < 0.05, **p < 0.01, ***p < 0.001; n = 4–8 rats per group.

Furthermore, AR-treated rats demonstrated superior physical performance, as evidenced by enhanced treadmill endurance **(Figure 1E)**. Histological analyses revealed a marked reduction in activated macrophages and fibroblasts—key contributors to myocardial fibrosis and calcification—in AR-treated hearts **(Figure 1F–M)**. This notable decrease in fibrotic and calcified myocardial areas, was accompanied by a better preservation of left ventricular wall structure and cardiomyocyte density **(Figure 2A, B)**.

**Figure 2.**
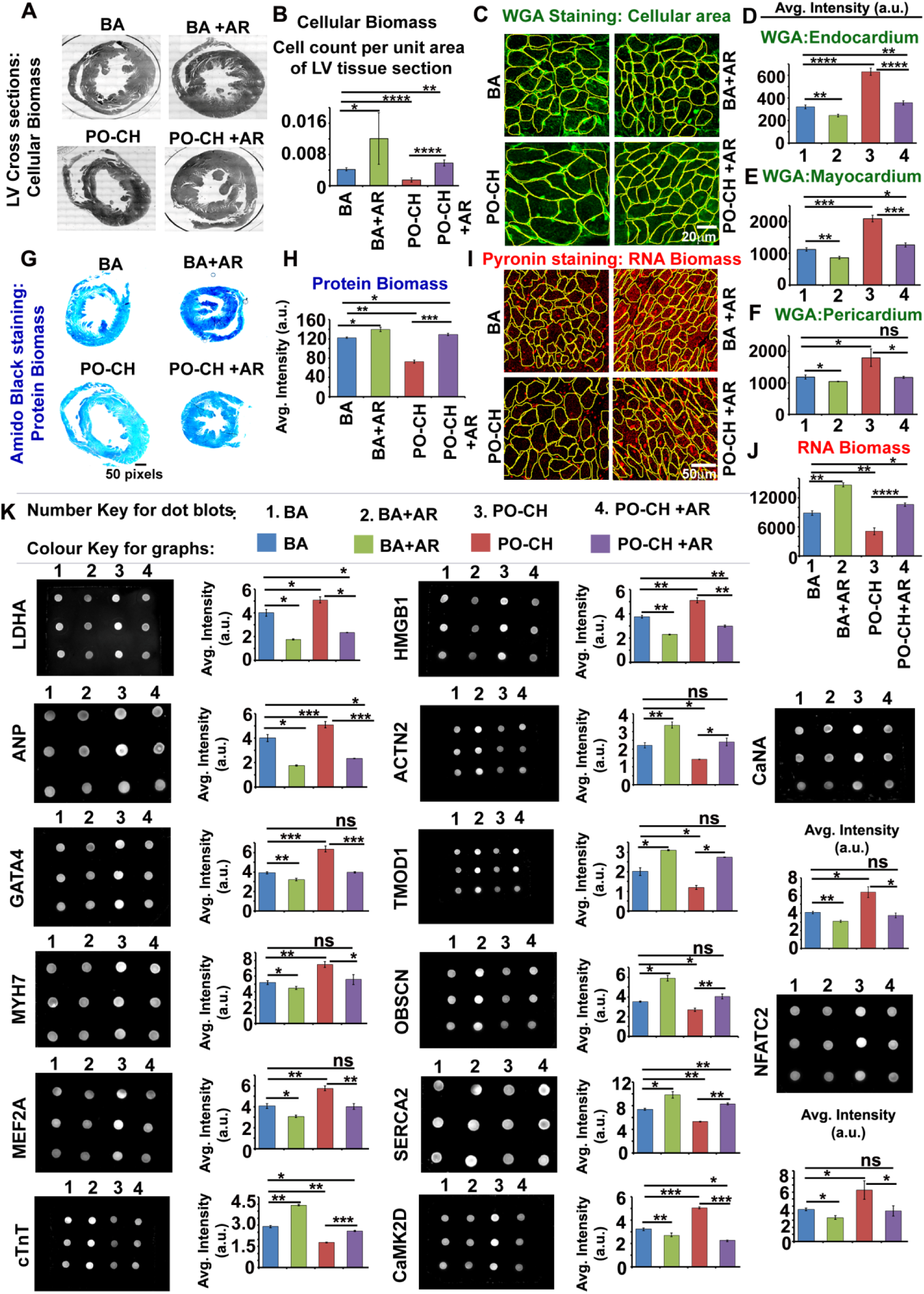
Additional validation of the Amalaki rasayana (AR)-based therapeutic model for left ventricular cardiac hypertrophy (CH). **(A)** Representative cross-sectional images of the left ventricular wall in 21-month-old PO-CH rats displaying dilated hypertrophic cardiomyopathy. **(B)** AR-treated BA and PO-CH rats exhibited reduced cellular loss and thicker ventricular walls compared to untreated counterparts, at the end of 21st month. **(C–F)** Wheat germ agglutinin (WGA) staining and analysis of cardiomyocyte surface area revealed decreased hypertrophy in AR-treated BA and PO-CH rats across endocardial, myocardial, and pericardial layers. **(G)** Although cardiomyocytes from untreated BA and PO-CH groups had larger surface areas, they showed reduced biomass, indicated by lower total protein (Amido Black staining) and RNA content (Pyronin Y staining) per cell, relative to AR-treated cohorts (measured over 5 random ROIs). **(K)** Serum levels of LDHA and HMGB1, markers of plasma membrane permeability and cell injury, were decreased in AR-treated groups, consistent with reduced cell loss as shown in panels A, B. Dot blot protein analysis demonstrated gene expression shifts in AR-treated rats: decreased fetal cardiac markers (ANP, GATA4, MYH7, MEF2A), increased adult cardiac muscle proteins (cTNT, ACTN2, TMOD1, OBSCN, SERCA2A), and reduced hypertrophy-associated proteins (CAMK2D, CaNA, NFATC2) compared to untreated BA and PO-CH groups. X-axis numerals in graphs correspond to: 1=BA, 2=BA+AR, 3=PO-CH, 4=PO-CH+AR. Data are mean ± SD; significance: *p < 0.05, **p < 0.01, ***p < 0.001; n = 8 rats/group. Image acquisition parameters were consistent across all conditions; single-cell analyses included 30–60 cells per independent experiment.

Morphometric analysis of cardiomyocytes showed a significant reduction in cell surface area in AR-treated rats, indicating attenuated myocyte hypertrophy **(Figure 2C–F)**. Additionally, there was an increase in total myocardial cellular biomass, suggesting improved tissue integrity and regenerative response **(Figure 2G–J)**. Biochemical assays revealed significantly lower serum levels of lactate dehydrogenase-A (LDHA) and high mobility group box 1 (HMGB1), indicating improved membrane integrity and reduced cellular damage in AR-treated rats **(Figure 2K, topmost panel)**.

Gene expression profiling further supported these findings. AR treatment was associated with significant downregulation of fetal and hypertrophy-associated genes (ANP, GATA4, MYH7, MEF2A, CAMK2D, CnNA, NFATC2) and upregulation of adult cardiac contractility markers (cTNT, ACTN2, TMOD1, OBSCN, SERC2A), suggesting a shift towards a physiologically mature myocardial phenotype **(Figure 2K, lower panels)**.

Given galectin-3’s well-established role in driving cardiac fibrosis, we hypothesized that Amalaki rasayana (AR) confers cardioprotective effects, at least in part, by attenuating the pathological signaling initiated by excessive surface-bound galectin-3. Notably, long-term AR administration (≥12 months) led to a significant decline in total serum oligomeric galectin-3 levels **(Figure 3A–D)**, with a pronounced reduction in the C-terminal galectin-3 epitope oligomers—particularly relevant in both BA and PO-CH models **(Figure 3E–G)**.

**Figure 3.**
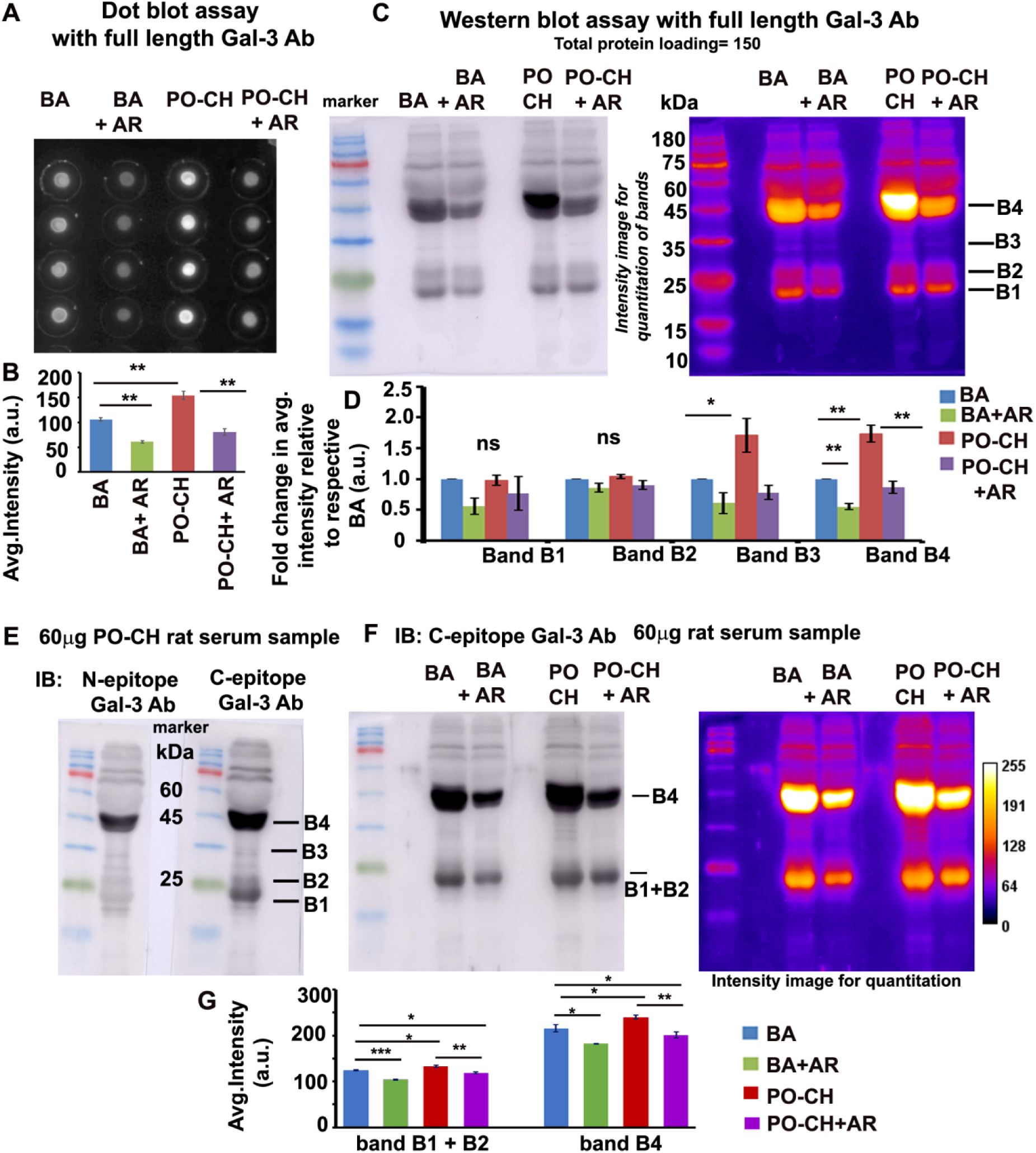
AR treatment reduces excessive galectin-3 C-epitope oligomer formation in cardiac hypertrophy. **(A,B)** Dot blot assay using a knockout-validated anti-galectin-3 antibody targeting full-length protein (aa 1–250) shows comparative galectin-3 levels in serum from BA, BA+AR, PO-CH, and PO-CH+AR groups (n = 4); each spot contains 6 µg serum. **(C, D)** Western blot quantification of full-length galectin-3 reveals differential monomer and oligomer enrichment across groups. **(E)** Analysis of 60 µg serum from PO-CH rats with epitope-specific antibodies indicates enrichment of C-epitope-containing galectin-3 in dimeric and oligomeric forms in the hypertrophy model. The N-epitope antibody recognizes N-N homo-dimers, homo-oligomers, and monomeric/oligomeric full-length protein; the C-epitope antibody recognizes C-C homo-dimers, homo-oligomers, and monomeric/oligomeric full-length protein. **(F, G)** Western blot validation confirms differences in monomeric and oligomeric C-epitope galectin-3 pools between AR-treated and untreated BA and PO-CH rats. Data are mean ± SD; significance by t-test: *p ≤ 0.05, **p ≤ 0.01, ***p ≤ 0.001.

Collectively, these results underscore the potential of AR as a promising therapeutic intervention for cardiac hypertrophy, with the galectin-3 C-epitope emerging as a critical molecular target. These findings warrant further investigation into galectin-3 C-epitope’s biomarker utility and translational relevance in human cardiac pathologies.

### The galectin-3 C-epitope oligomers play a dominant role in the generation of cardiac hypertrophy

To elucidate the role of galectin-3 C-epitope oligomers in the pathogenesis of cardiac hypertrophy, we employed a molecular strategy involving the expression of EGFP-tagged full-length wild-type galectin-3 (Gal-3 WT-EGFP) and a carbohydrate recognition domain mutant (Gal-3 R186S, hereafter referred to as CRD-MUT-Gal-3 EGFP), which is deficient in binding β-galactoside-containing carbohydrate moieties [25]. HEK293 cells were used as a heterologous overexpression system to ensure high transfection efficiency (∼100%).

To simulate hypertrophic stress, we subjected one set of cells to pressure overload (PO) using an established mechanical weight application method [26, 27], while another set served as an untreated control (No PO). After 24 hours, conditioned media were collected from both Gal-3 WT-EGFP and CRD-MUT-Gal-3 EGFP expressing cells. Fluorescence intensities of the secreted fusion proteins were measured using a multimode plate reader, and media were normalized to equal fluorescence intensity to ensure consistent galectin-3 protein levels across samples (2.5 mL per condition; see experimental schema, **Figure 4A**).

**Figure 4.**
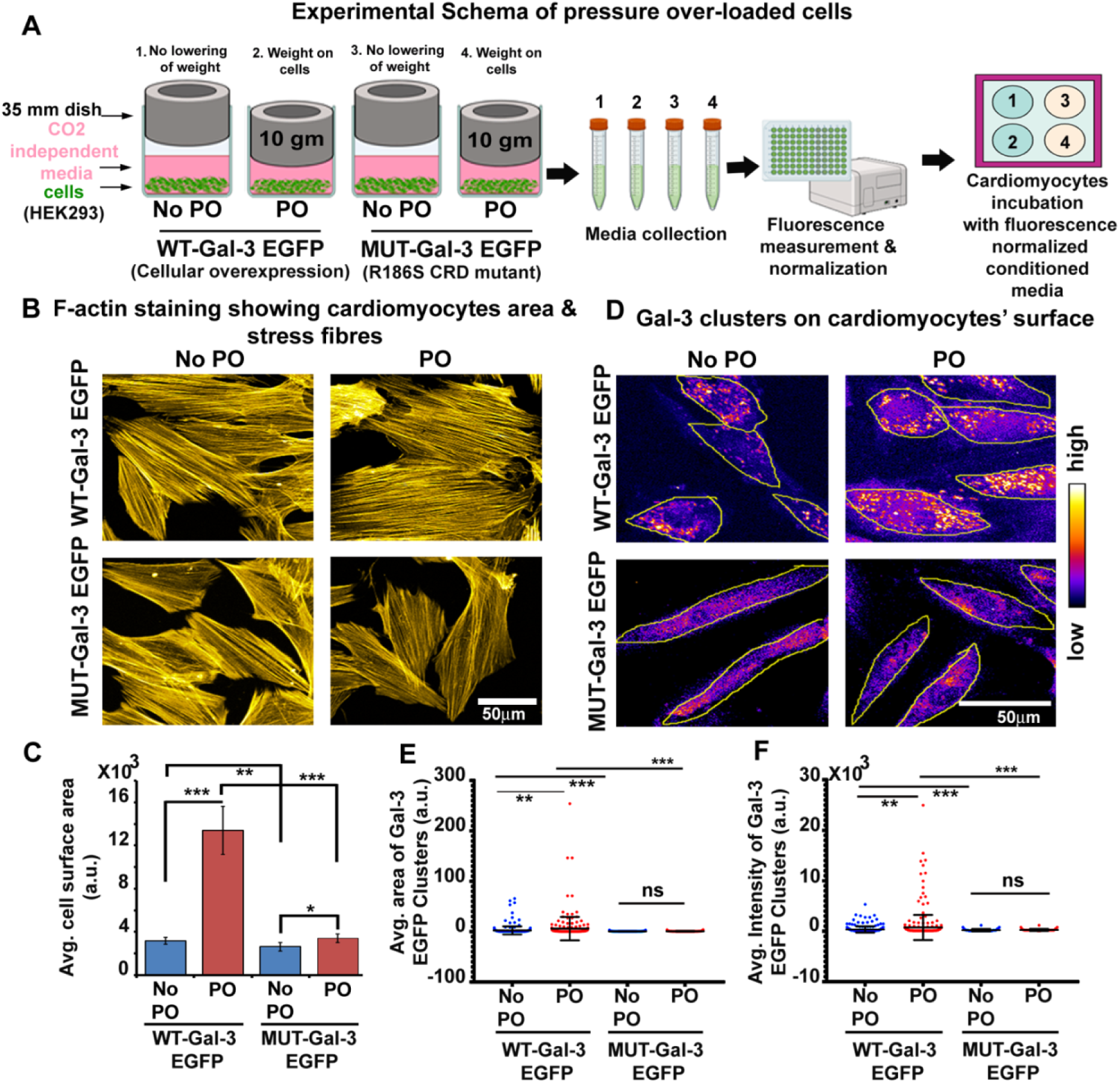
Predominance of galectin-3 C-epitope oligomers in cardiac hypertrophy. **(A)** Schematic of the in vitro cellular pressure overload (PO) model, created with Biorender.com **(B, C)** Representative rhodamine phalloidin-stained images and quantification of cardiomyocyte surface area following incubation with galectin-3 WT-EGFP or carbohydrate-binding mutant (Gal3 R186S, CRD-MUT) under no pressure overload (No-PO) and PO conditions. **(D–F)** Analyses of binding patterns, signal intensity, and cluster area of surface-bound WT-EGFP and CRD-MUT-EGFP galectin-3 on cardiomyocytes, under No-PO and PO conditions. Data are mean ± SD; significance by t-test: *p ≤ 0.05, **p ≤ 0.01, ***p ≤ 0.001.

The fluorescence-normalized conditioned media were then applied to primary cardiomyocytes for 10 hours, followed by cell fixation and morphometric analysis. Quantification of cardiomyocyte surface area revealed significantly less hypertrophic enlargement in cells treated with CRD-MUT-Gal-3 EGFP media compared to those exposed to Gal-3 WT-EGFP, under both No PO and PO conditions **(Figure 4B, C)**. Fluorescence imaging further indicated that the CRD-MUT-Gal-3 exhibited markedly reduced surface binding, correlating with diminished hypertrophic response **(Figure 4D, F)**.

Notably, Gal-3 WT-EGFP derived from PO-conditioned media displayed a distinct micron-sized clustered pattern on the cardiomyocyte surface. In contrast, CRD-MUT-Gal-3 EGFP exhibited a more diffuse and uniform staining pattern, suggesting a key role of the C-epitope in oligomerization and membrane clustering **(Figure 4D,E)**.

To examine whether Amalaki rasayana (AR) modulates galectin-3 secretion and cell surface binding to prevent hypertrophy, we subjected cardiomyocytes to 24-hour PO stress followed by treatment with 100 µg/mL AR (dose and viability determined from MTT and cell count assays respectively; **Figure 5A, B)** for an additional 24 hours. The supernatant (media) analysis revealed significantly decreased levels of unbound galectin-3 C-epitope oligomers in AR-treated groups under both No PO and PO conditions, compared to their respective untreated controls **(Figure 5C, D)**. Similarly, surface-bound C-epitope signal intensity was markedly reduced upon AR treatment **(Figure 5E, F)**.

**Figure 5.**
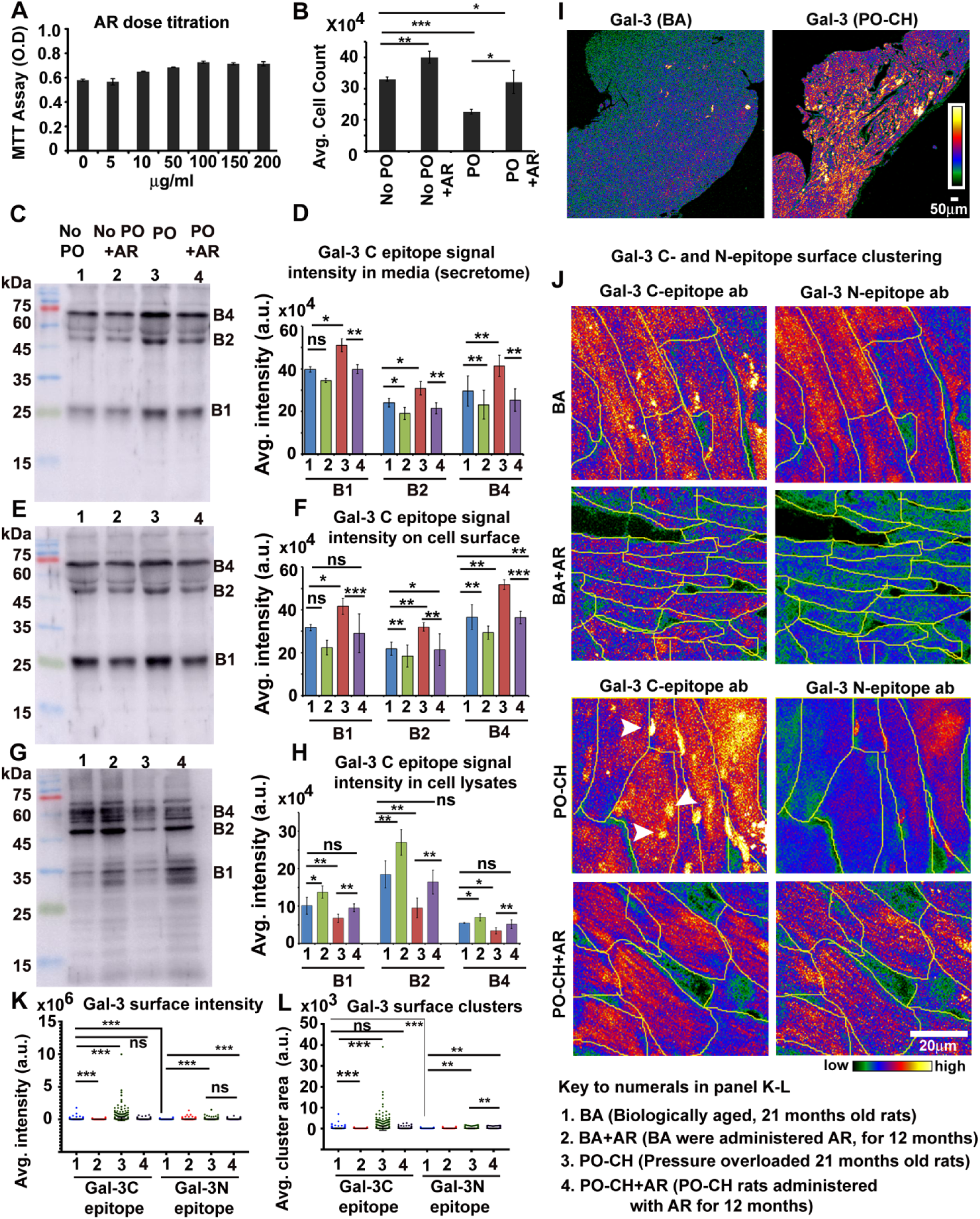
AR treatment significantly reduces extracellular galectin-3 C-epitope oligomers in vitro and in vivo. **(A)** MTT assay assessing cardiomyocyte viability with increasing AR doses over 24 hours; no viability increase observed above 100 µg/ml. **(B)** Cardiomyocytes subjected to 24-hour pressure overload (PO) were treated with 100 µg/ml AR for an additional 24 hours; AR treatment significantly increased cell counts compared to No-PO and PO alone. **(C–H)** Western blot analysis quantifying monomeric and oligomeric galectin-3 C-epitope levels in culture media (C, D), surface-bound fractions (E, F), and intracellular lysates (G, H) under conditions described in (B). Surface galectin-3 was isolated via low pH wash. **(I)** Representative 4× images of surface-bound galectin-3 C-epitope clusters in left ventricular tissues from BA and PO-CH rats, showing high clustering in PO-CH. **(J–L)** Comparative quantification of galectin-3 N- and C-epitope surface levels in BA and PO-CH tissues with or without AR treatment; untreated PO-CH tissues exhibit significant C-epitope clustering. Graph and blot labelling for C-H: 1=No PO, 2=No PO+AR, 3=PO, 4=PO+AR. Graph labelling for K-L: 1=BA, 2=BA+AR, 3=PO-CH, 4=PO-CH+AR. Data are mean ± SD; n ≥ 4 animals/group; significance by t-test: *p ≤ 0.05, **p ≤ 0.01, ***p ≤ 0.001.

Interestingly, Western blot analysis of cell lysates revealed a marked increase in intracellular galectin-3 levels in AR-treated groups **(Figure 5G, H)**, suggesting that AR may hinder excessive galectin-3 secretion by altering its intracellular trafficking or release mechanisms [28, 29].

These findings were further validated in an in vivo rat model. Cardiomyocytes from hypertrophied left ventricles under biological ageing (BA) and pressure overload-induced cardiac hypertrophy (PO-CH) exhibited high levels of membrane-bound C-epitope oligomers, which were significantly reduced in their respective AR-treated counterparts **(Figure 5I–L)**. Moreover, surface staining in PO-CH tissues revealed intense C-epitope clustering that exceeded the N-epitope signal, supporting a dominant surface residency of C-terminal galectin-3 oligomers in the hypertrophic state **(Figure 5J–L)**.

Finally, human cardiac hypertrophy (CH) tissue samples exhibited surface staining patterns for both N- and C-terminal epitopes that closely mirrored those observed in the rat model, reinforcing the translational significance of these findings **(Figure 6A, B**).

**Figure 6.**
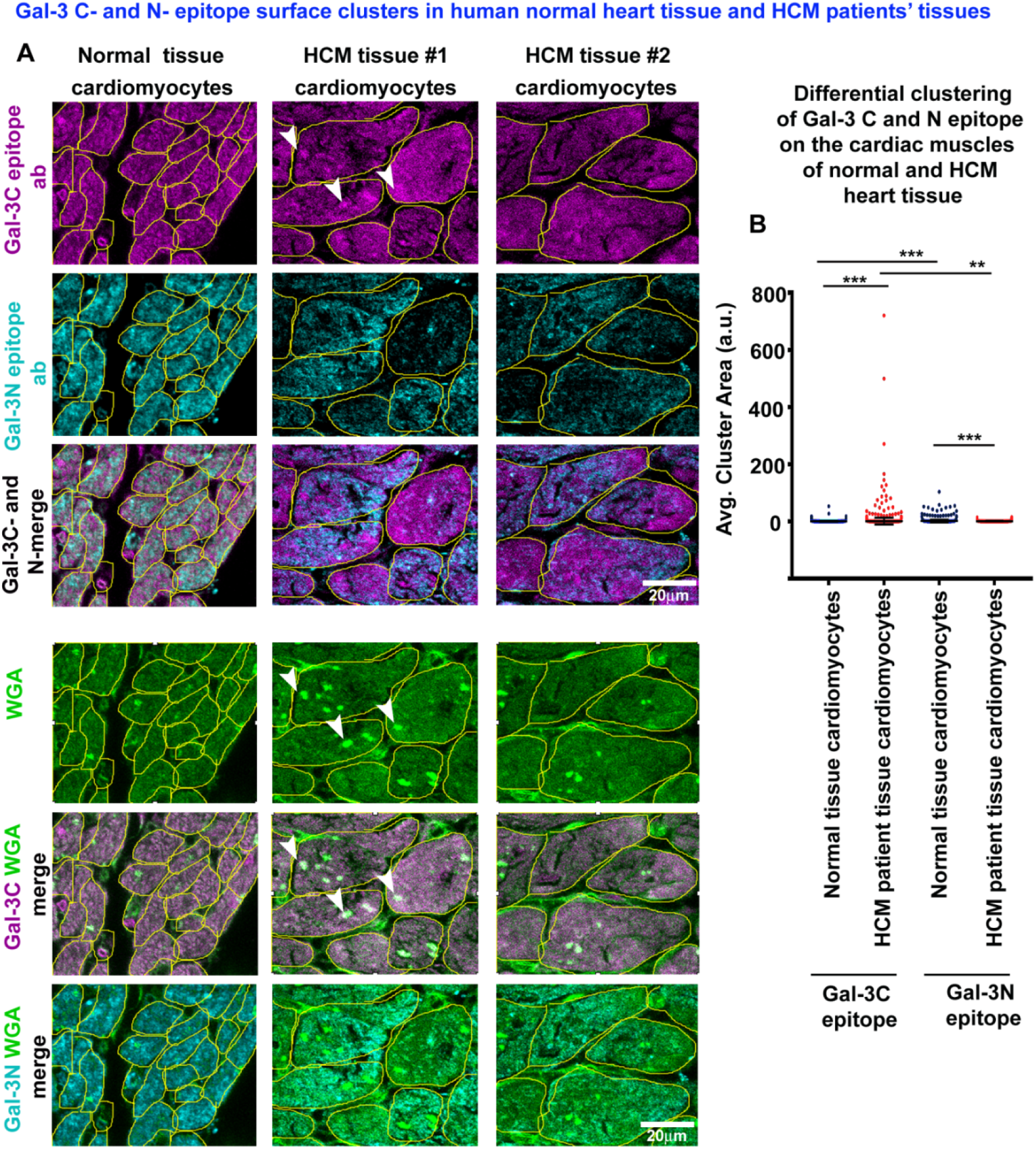
Validation of elevated extracellular galectin-3 C-epitope oligomers in hypertrophic cardiomyopathy (HCM) donor cardiomyocytes versus normal controls. **(A,B)** Human cardiac tissues from normal and HCM donors were stained for surface-bound galectin-3 N-epitope (cyan) and C-epitope (magenta). Merged images reveal significantly reduced N and C-epitope co-localization in HCM cardiomyocytes compared to normal tissue, indicating predominance of C-C homo-oligomers over N-N or full-length galectin-3. Wheat germ agglutinin (WGA, green) marks cardiomyocyte surfaces. Merged images of C-epitope and WGA highlight prominent galectin-3 C-epitope clusters (white arrowheads) on HCM cardiomyocytes. Data are mean ± SD; significance by t-test: *p ≤ 0.05, **p ≤ 0.01, ***p ≤ 0.001.

### **G**allic acid (GA) in AR extract effectively reduces excess extracellular levels of galectin-3 C-epitope oligomers

To identify bioactive constituents within Amalaki rasayana (AR) responsible for the observed reduction in extracellular galectin-3 C-epitope (Gal-3C) oligomers, we performed in silico docking and molecular dynamics (MD) simulations with 18 phytocompounds previously identified in AR [18]. Among them, three compounds displayed strong binding affinities toward the Gal-3C domain, with gallic acid (GA) emerging as a prominent candidate **(Figure 7A)**.

**Figure 7.**
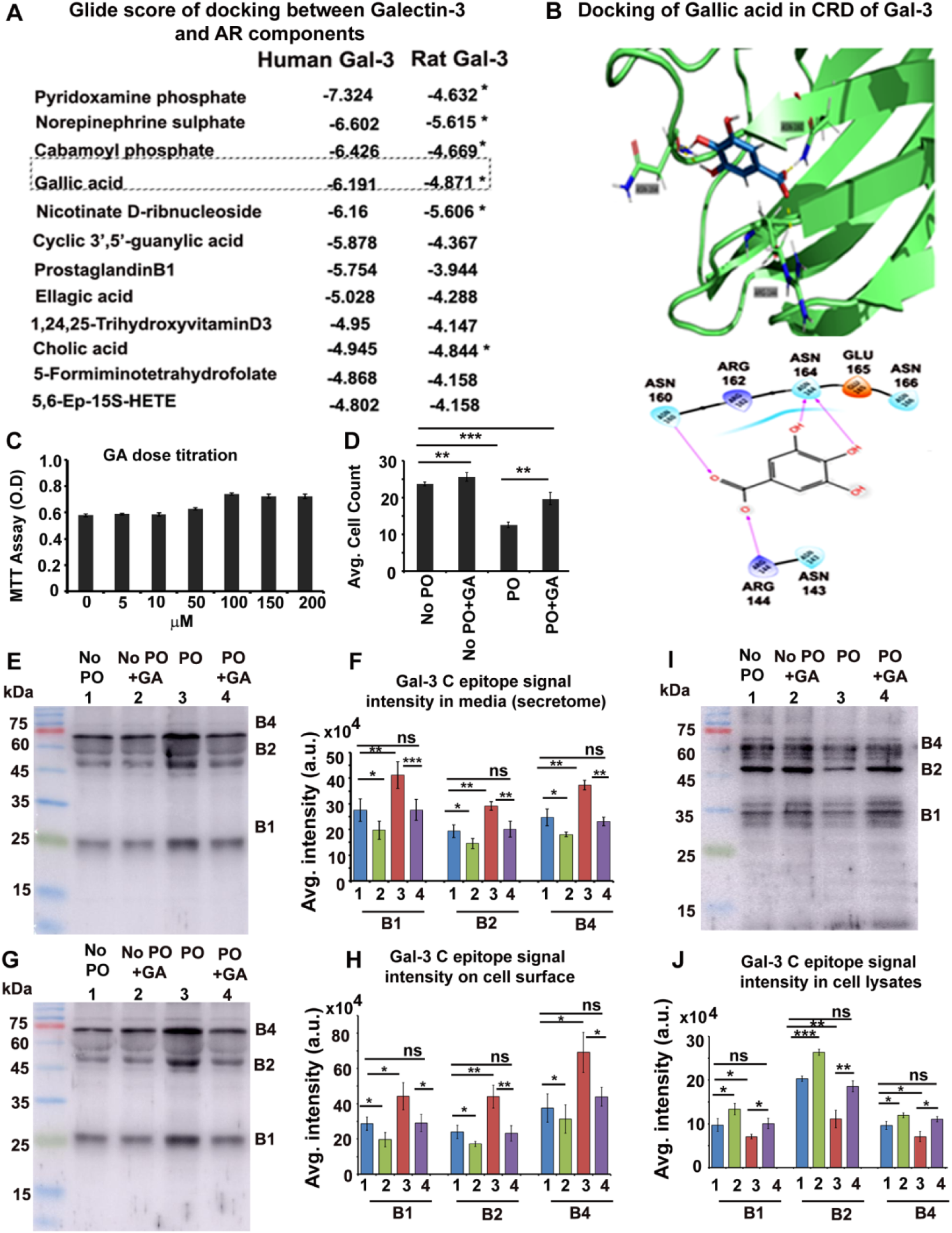
Gallic acid (GA) from AR extract reduces excess extracellular galectin-3 C-epitope oligomers. **(A)** Glide docking scores of 18 key AR components with rat and human galectin-3. **(B)** Molecular dynamics simulation showing GA forming hydrogen bonds with three critical residues (Arg144, Asn160, Asn164) in the galectin-3 carbohydrate recognition domain (CRD). **(C)** MTT assay assessing cardiomyocyte viability with increasing GA concentrations over 24 hours; no significant viability increase above 100 µM GA. **(D)** Cardiomyocytes subjected to 24-hour pressure overload (PO) were treated with 100 µM GA for an additional 24 hours; GA treatment significantly increased cell counts compared to No-PO and PO alone. **(E–J)** Western blot analysis quantifying monomeric and oligomeric galectin-3 C-epitope levels in culture media (E, F), surface-bound fractions (G, H), and intracellular lysates (I, J) under conditions described in (D). Surface galectin-3 was isolated via low pH wash. Graph and blot labelling for E-J: 1=No PO, 2=No PO+AR, 3=PO, 4=PO+AR. Data are mean ± SD; significance by t-test: *p ≤ 0.05, **p ≤ 0.01, ***p ≤ 0.001.

Given AR’s enrichment in GA, we further investigated GA’s role in modulating Gal-3C oligomerization and glycan-binding properties. MD simulations revealed that GA binds to the carbohydrate recognition domain (CRD) of Gal-3C via hydrogen bonding interactions with key residues Arg144, Asn160, and Asn164 **(Figure 7B)**, suggesting a potential mechanism for its modulatory action.

Functional assays on cardiomyocytes treated with 100 µM GA—an effective concentration confirmed by MTT and cell count analyses **(Figure 7C, D)—**revealed a significant decrease in extracellular Gal-3C oligomers in both the media and cell surface-bound fractions.

Conversely, cell lysates exhibited an increased galectin-3 signal, indicative of enhanced intracellular retention **(Figure 7E–J)**. This pattern closely parallels the effects observed with AR, reinforcing GA as a key active component.

Cellular imaging further confirmed that Gal-3C epitope signals were more intense and clustered than those of the N-terminal epitope in untreated cells. Upon treatment with GA or AR, both signal intensity and clustering were significantly diminished in pressure overload (PO) and control (No PO) conditions **(Figure 8A–C)**. These effects were comparable to those observed with BAPTA-AM, a calcium chelator, suggesting that GA and AR may reduce galectin-3 secretion by lowering intracellular calcium levels—an established trigger of galectin-3 exocytosis [28].

**Figure 8.**
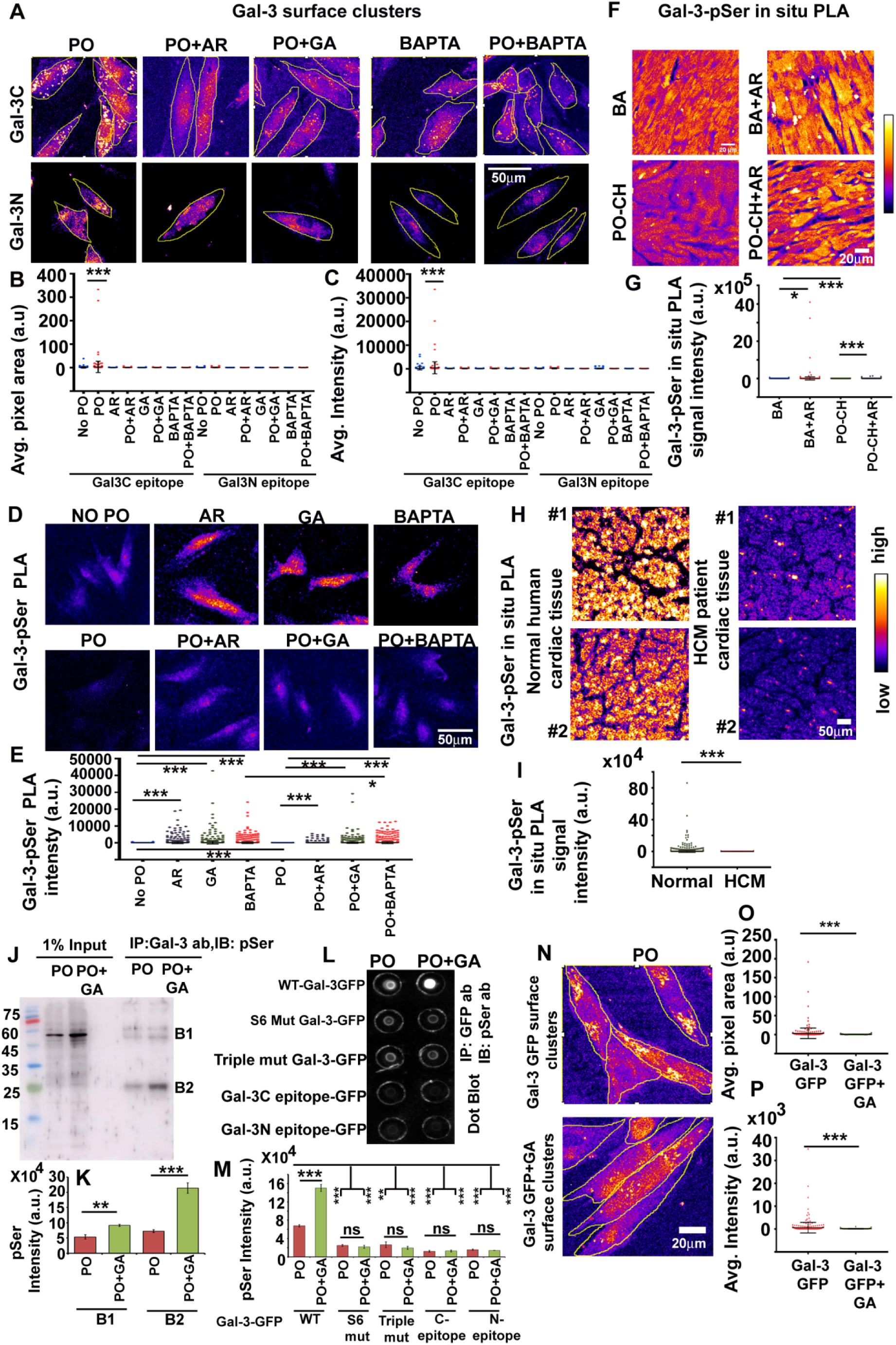
Gallic acid (GA) in AR reduces galectin-3 release by promoting N-epitope phosphorylation and cytoplasmic retention. **(A–C)** GA, AR, and BAPTA-AM treatments reduce extracellular galectin-3 C-epitope oligomer cluster area and intensity on cardiomyocyte surfaces under pressure overload (PO) and No-PO conditions. **(D, E)** Proximity ligation assay (PLA) shows increased galectin-3 phosphorylation and intracellular retention in AR- and GA-treated cardiomyocytes. BAPTA-AM–mediated calcium reduction enhances phosphorylation and retention, suggesting calcium regulates galectin-3 secretion and intracellular loss. **(F, G)** PLA quantification of phosphorylated galectin-3 in BA and PO-CH rat tissues with or without AR treatment; untreated PO-CH tissues show significantly lower phosphorylation. **(H, I)** PLA analysis comparing normal human left ventricular and hypertrophic cardiomyopathy (HCM) tissues reveals reduced phosphorylated galectin-3 in HCM cardiomyocytes. **(J, K)** Immunoprecipitation (IP) with anti-galectin-3 and western blotting mediated immuno-detection with pan phosphoserine antibodies demonstrates enhanced intracellular galectin-3 phosphorylation in GA-treated cardiomyocytes. **(L, M)** IP of GFP-tagged WT galectin-3 from PO ± GA cardiomyocytes shows increased phosphorylation with GA; no change is observed with S6 mutant (that inhibits serine phosphorylation at 6^th^ position in N-epitope), and in the triple mutant that inhibits GA-binding in the CRD, indicating GA binds galectin-3 CRD and modulates phosphorylation at N-epitope S6. No change in phosphorylation is observed in IP samples expressing either only C- or N-epitope, again suggesting that GA influences N-epitope serine phosphorylation by binding to the C-epitope. **(N–P)** Incubation of fresh cardiomyocytes with GA-pre-treated media from WT-GFP galectin-3–expressing PO cardiomyocytes shows reduced surface binding GFP-galectin-3 in a GA dependent manner, indicating GA inhibits surface binding of extracellular galectin-3. Data are mean ± SD; n ≥ 4 animals or 3 independent experiments; significance by t-test: *p ≤ 0.05, **p ≤ 0.01, ***p ≤ 0.001.

To explore intracellular retention mechanisms, we performed in situ proximity ligation assays (PLA), which demonstrated that GA enhances cytoplasmic retention of Gal-3, likely by promoting serine phosphorylation at the S6 residue of the N-terminal domain—a modification catalyzed by casein kinase 1 (CKI) **(Figure 8D, E)** [29, 30]. Consistent with calcium regulation, BAPTA-AM also increased Gal-3 S6 phosphorylation and retention, indicating that reduced cytosolic calcium facilitates CKI-mediated phosphorylation.

Supporting in vitro findings, left ventricular tissues from both biological ageing (BA) and PO-induced hypertrophy (PO-CH) animal models showed significantly higher phosphorylated Gal-3 levels in AR-treated cohorts **(Figure 8F, G)**. Furthermore, cardiac tissues from hypertrophic cardiomyopathy (HCM) treatment naive patients exhibited markedly lower Gal-3 phosphorylation than healthy donor hearts, consistent with excessive extracellular release and decreased cytoplasmic retention in pathological states **(Figure 8H, I)**.

Immunoprecipitation (IP) of cardiomyocyte lysates using an anti-Gal-3 antibody revealed that GA-treated cells retained a highly phosphorylated intracellular Gal-3 pool compared to untreated controls **(Figure 8J, K)**. To dissect the specificity of this GA-induced phosphorylation, we engineered Gal-3 carbohydrate recognition domain (CRD) mutants lacking GA-binding sites and overexpressed them in cardiomyocytes. Upon exposure to pressure overload (PO) with or without GA, cells expressing triple CRD mutants exhibited markedly reduced Gal-3 phosphorylation compared to wild-type Gal-3 **(Figure 8L, M)**, indicating that GA-binding sites are essential for phosphorylation.

Moreover, cells expressing the S6A Gal-3 mutant—unable to undergo CKI-mediated phosphorylation at the N-epitope—showed no phosphorylation response to GA treatment, confirming that serine-6 is a critical site for GA-driven phosphorylation **(Figure 8L, M)**. Consistently, IP samples expressing only the C- or N-epitope alone displayed no detectable phosphorylation, further supporting the model that GA modulates N-terminal serine phosphorylation indirectly through its interaction with the C-epitope **(Figure 8L, M)**.

To assess the likely inhibitory impact of GA on excess extracellular galectin-3 pool, we pre-mixed PO-conditioned media from WT Gal-3-GFP-expressing cardiomyocytes with GA (100µM) before incubation on fresh cardiomyocytes. Surface fluorescence analysis revealed significantly reduced Gal-3 binding in a GA-dependent manner **(Figure 8N–P)**, suggesting that GA interferes with surface interactions of Gal-3.

Collectively, these data suggest that GA acts through dual mechanisms: (1) by chelating cytosolic calcium, thus reducing secretion of Gal-3C oligomers, and (2) by binding to the Gal-3 CRD, inducing conformational changes that expose the S6 site for CKI-mediated phosphorylation. This dual action promotes intracellular retention and reduces surface clustering of galectin-3, thereby mitigating its hypertrophic effects.

Indeed, literature supports this mechanistic framework. Elevated cytosolic calcium—common under mechanical stress or pressure overload—activates calcineurin A, a phosphatase that antagonizes CKI activity, thereby reducing galectin-3 phosphorylation and enhancing its secretion [31, 32]. Our findings suggest that GA, by modulating both calcium signaling and phosphorylation, shifts galectin-3 toward a non-pathogenic intracellular fate, offering a promising therapeutic strategy.

Ongoing studies aim to further delineate the structural basis by which GA and AR influence galectin-3 conformations, phosphorylation dynamics, and calcium homeostasis in cardiac hypertrophy.

### Sera and tissue samples from human donors confirm the pivotal role of the galectin-3 C-epitope in the generation of cardiac hypertrophy (CH)

Literature suggests that during cardiac hypertrophy (CH), the plasma membrane of cardiomyocytes experiences dynamic changes due to hydrostatic pressure. This mechanical load and accompanying membrane stretch are sensed by surface mechanosensors, which activate the RhoA-ROCK1 signaling pathway to stabilize F-actin stress fibers that help resist the mechanical strain [33]. During transient pressure, F-actin stress fibers serve a protective role; however, to accommodate extended fiber length, hypertrophy-associated genes are upregulated, leading to increased cardiomyocyte surface area.

Under sustained mechanical load, however, the RhoA-ROCK1 axis and extensive stress fiber formation lead to maladaptive signaling. RhoA-induced cytoplasmic calcium elevation triggers aberrant activation of hypertrophy-associated genes such as NFATC2, MEF2A, and GATA4 [33, 34]. Rigid, bundled F-actin stress fibers deform the nucleus, affecting chromatin stability, and simultaneously cause disassembly of caveolae—the principal mechanoprotective membrane invaginations—by removing the caveolar coat protein PTRF. This leads to heightened membrane tension [35].

Elevated cytosolic calcium also promotes the over-secretion of galectin-3 [28]. The secreted galectin-3 binds and activates membrane receptors such as EGFR, CD44, and integrins, initiating their autophosphorylation [36, 37, 38, 39]. This further amplifies deleterious RhoA-ROCK1 signaling and activates cSrc kinase, which phosphorylates CAV1, a structural caveolae protein, resulting in additional caveolar loss [40, 41]. This cascade accelerates CH progression, increases membrane permeability, and contributes to cardiomyocyte loss. Excess circulating galectin-3 may also act systemically, leading to dysfunction in peripheral tissues.

Therefore, to evaluate the clinical relevance of galectin-3 C-epitope in human hypertrophic cardiomyopathy (HCM), we analyzed serum samples from young and old healthy donors and HCM survivors undergoing treatment. Atrial natriuretic peptide (ANP), secreted by cardiomyocytes in response to mechanical stress, served as a tension biomarker **(Figure 9A, B)**. Older individuals and HCM patients had elevated ANP levels compared to younger donors, reflecting increased myocardial wall stress. Notably, ANP levels in HCM patients were comparable to aged healthy controls, suggesting therapeutic benefit from clinical interventions.

**Figure 9.**
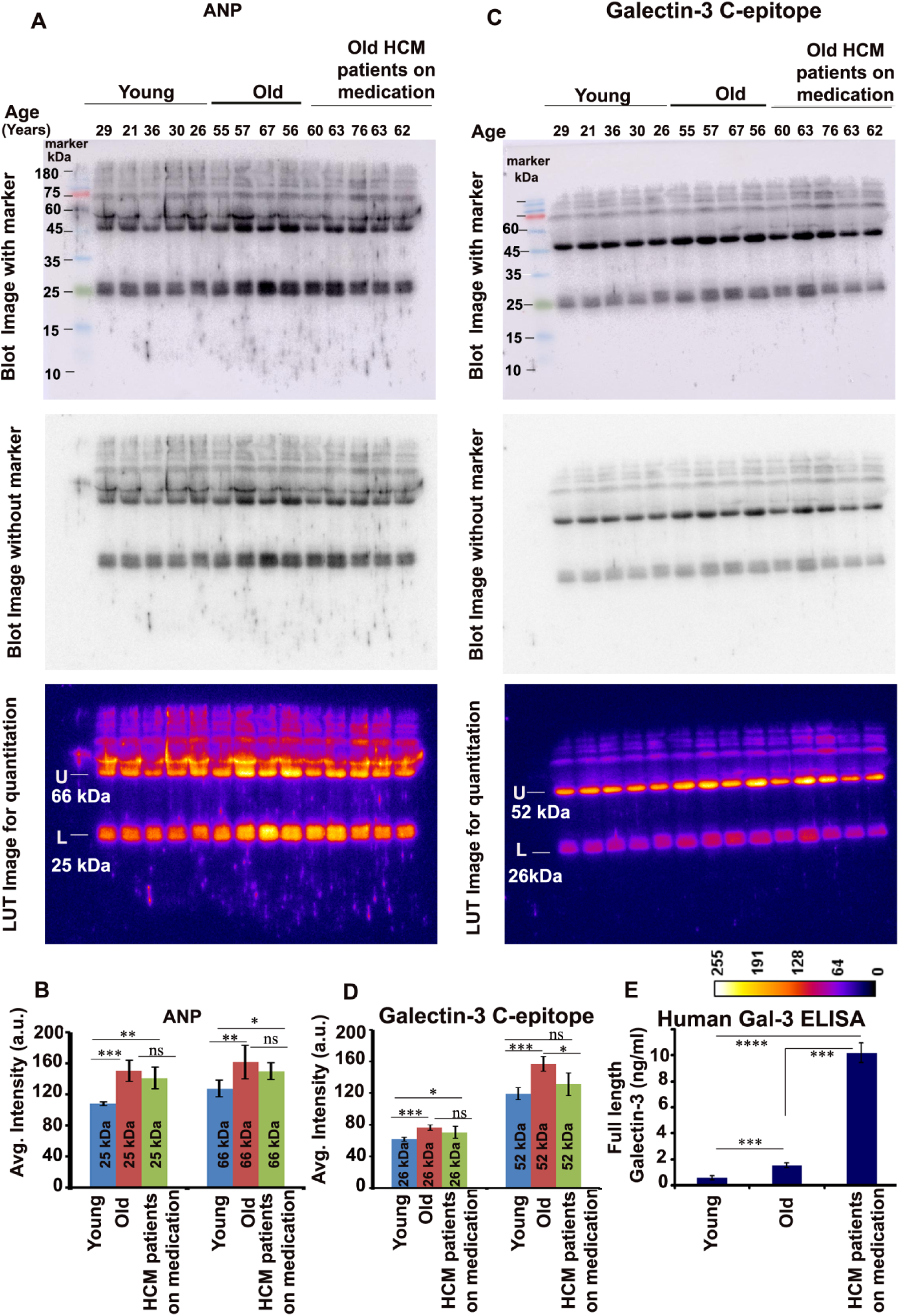
Validation of galectin-3 C-epitope oligomers in human donor serum samples. **(A–D)** Serum from young individuals, older adults, and cardiac hypertrophy survivors (post various pharmacological treatments) was analyzed for ANP and galectin-3 C-epitope levels. **(E)** Galectin-3 concentrations in these sera were quantified by a commercial ELISA kit, wherein the plates were coated using a full-length anti-galectin-3 antibody. Statistical significance was assessed by t-tests: *p ≤ 0.05, **p ≤ 0.01, ***p ≤ 0.001.

We next analyzed serum galectin-3 C-epitope oligomers via Western blotting **(Figure 9C, D)**. HCM survivors exhibited slightly lower C-epitope oligomer levels than aged healthy controls, implying reduced cardiac strain due to treatment. Young donors with low ANP also showed lower C-epitope levels. Interestingly, ELISA analysis for total galectin-3 did not correlate with the oligomer profile; HCM survivors showed the highest total galectin-3 levels among all groups **(Figure 9E)**. This discrepancy underscores the limitations of ELISAs using full-length galectin-3 antibodies, which may not adequately detect C-epitope oligomers, thereby obscuring galectin-3’s potential as a biomarker of treatment response in CH.

Although drug-naïve HCM patient sera were unavailable, our tissue analysis has revealed significantly higher C-epitope oligomer accumulation on cardiomyocyte surfaces in HCM drug-naïve hearts compared to age-matched controls **(Figure 6)**. Thus, reduced serum C-epitope levels in treated HCM patients likely reflect therapeutic efficacy, highlighting the utility of C-epitope oligomers as a treatment response biomarker.

Given that serum reflects tissue-level metabolic and secretory states, we investigated how donor sera affected cardiomyocyte hypertrophy. Cardiomyocytes incubated with sera from older donors exhibited increased cell surface area compared to those treated with sera from HCM patients on medication **(Figure 10A, B**), mirroring the C-epitope oligomer profiles (as shown in Figure 9C-D). To directly assess the role of the C-epitope, we enriched aged donor serum fractions for N- and C-epitopes via immunoprecipitation. Cardiomyocytes exposed to C-epitope-enriched fractions displayed significantly more hypertrophy than those treated with N-epitope-enriched fractions **(Figure 10C, D)**. Pre-incubation of the C-epitope fraction with lactose—a glycan-binding blocker—significantly attenuated this hypertrophy **(Figure 10C, D)**.

**Figure 10.**
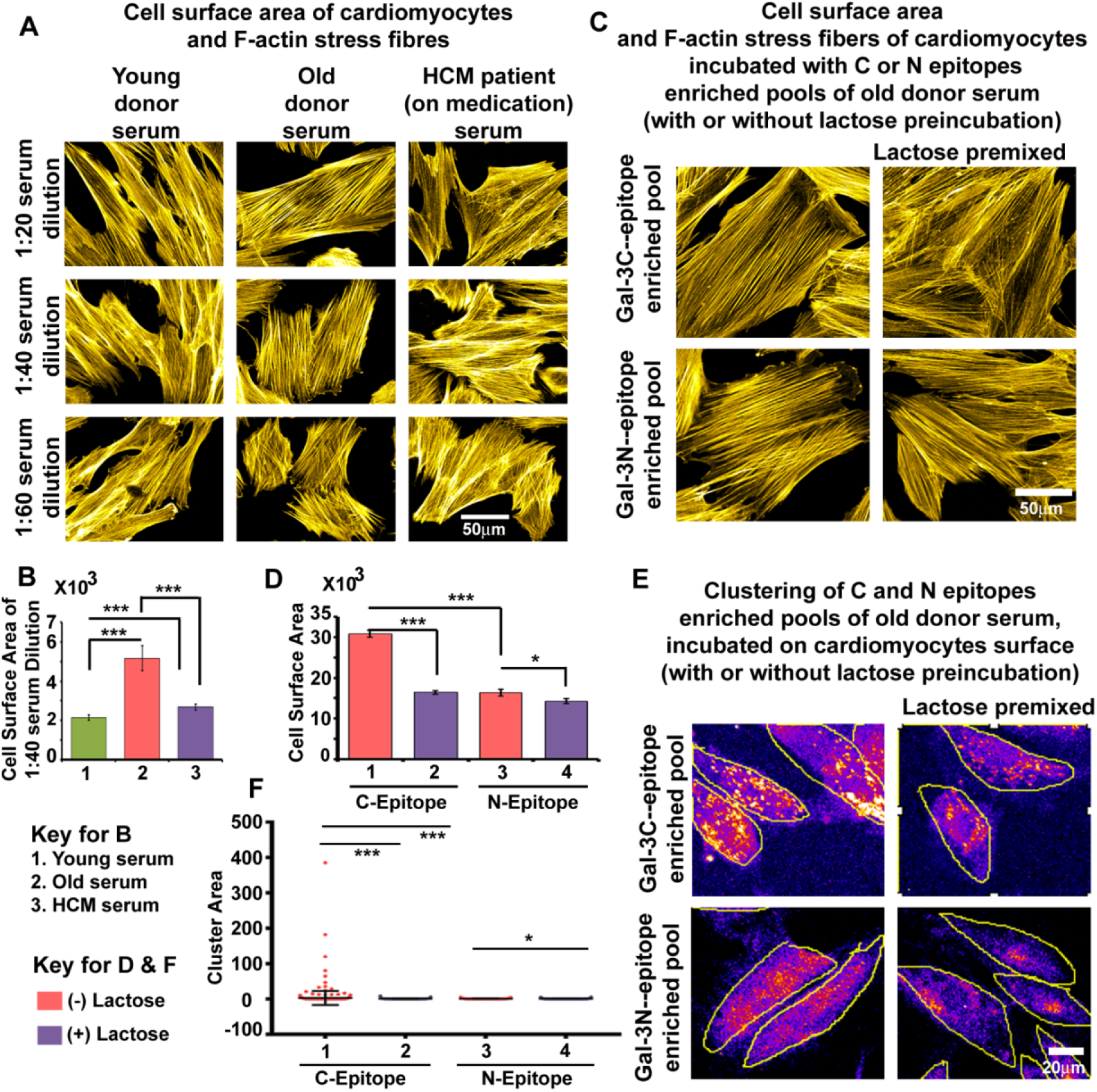
Validation of galectin-3 C-epitope oligomer dominance in cardiomyocyte hypertrophy over N-epitope. **(A,B)** Rat cardiomyocytes treated with sera from young, old, and cardiac hypertrophy survivors at varying dilutions were stained with rhodamine-phalloidin to assess cell surface area and visualize stress fibers, which increase under mechanical stress. **(C, D)** Galectin-3 C- and N-epitope pools were immunoprecipitated from human old donor sera. Pre-incubation with lactose or vehicle was performed before conditioning cardiomyocytes with these pools. Changes in cell surface area and stress fiber formation were quantified, showing C-epitope-enriched sera induced hypertrophy, which lactose attenuated. **(E, F)** Immunofluorescence analysis revealed differential cell surface binding patterns of Gal- 3 C- and N-epitope pools, with C-epitope exhibiting clustered organization and inducing prominent stress fibers, modulated by lactose treatment. Keys to histogram bar colours and numericals used on the X-axis of the graphs are provided within the figure. Statistical significance was evaluated by t-tests: *p ≤ 0.05, **p ≤ 0.01, ***p ≤ 0.001.

Furthermore, the surface clustering of C-epitope oligomers was markedly higher than that of N-epitopes, unless blocked by lactose **(Figure 10E, F)**. These results suggest that the galectin-3 C-epitope induces hypertrophy through its carbohydrate recognition domain (CRD), which facilitates oligomer binding to cell surfaces and activates mechanotransductive stress pathways.

Treatment of cardiomyocytes with gallic acid (GA) mimicked and exceeded the inhibitory effects of lactose on C-epitope-induced hypertrophy and surface oligomerization **(Figure 11A–D)**. In comparison to the N-epitope, C-epitope administration also elevated RhoA and ROCK1 expression **(Figure 11E–G)** [33, 34], enhanced F-actin stress fiber formation **(Figure 11C)** [42], promoted caveolae disassembly **(Figure 11H, I)** [40, 41], and increased cellular tension (increase in nuclear YAP1, **Figure 11J, K)** [43]. It also elevated nuclear envelope tension (increase in nuclear Lamin A, **Figure 11L, M**) [44], induced DNA damage **(Figure 11N, O)**, and upregulated hypertrophic gene expression **(Figure 11P–S)** [33]. Additionally, increased membrane permeability and cell loss were observed **(Figure 11T, U)**. These pathological effects were significantly mitigated by lactose and GA, both of which block the CRD of galectin-3.

**Figure 11.**
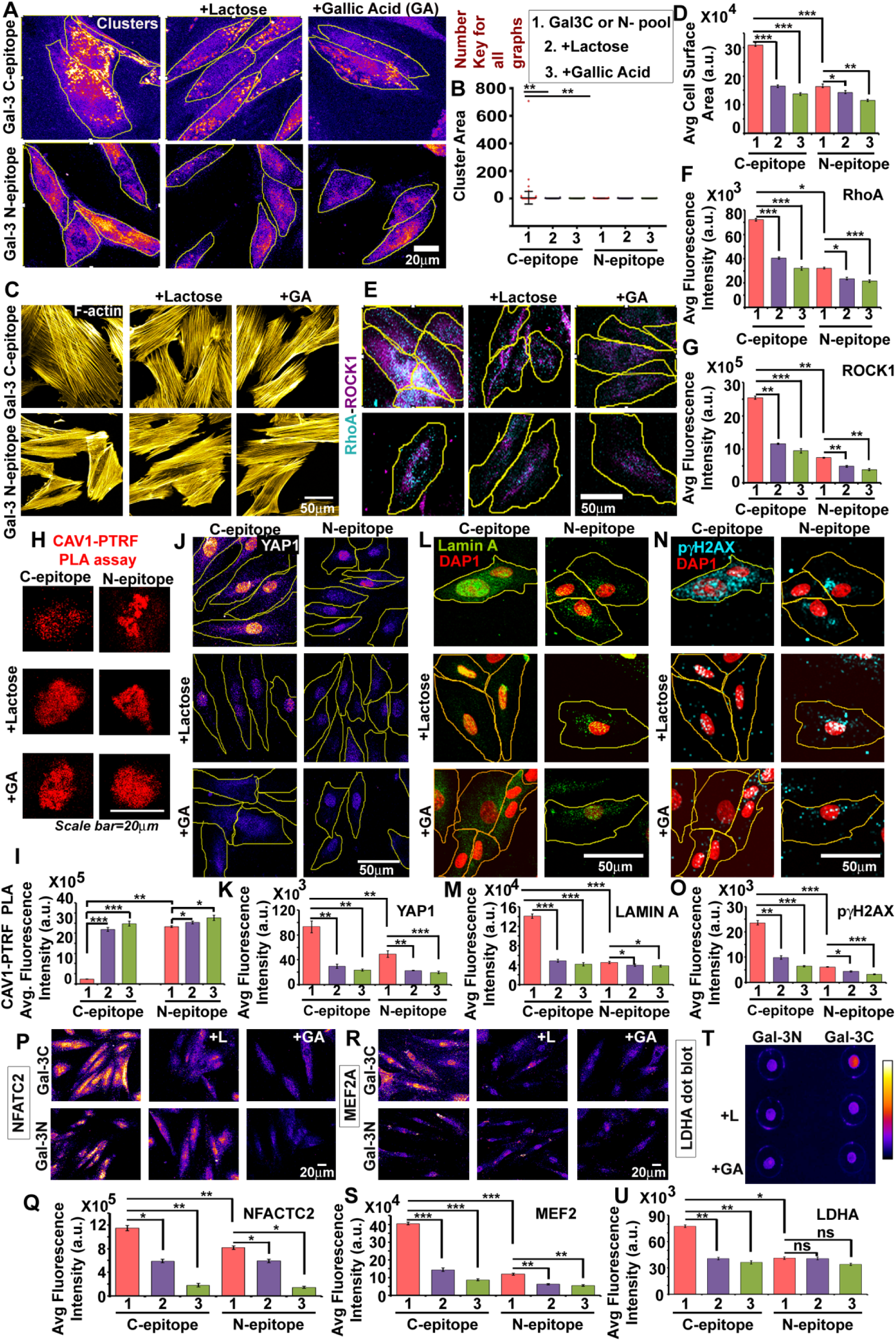
Human galectin-3 C-epitope oligomers play a dominant role in cardiac hypertrophy and cardiomyocyte loss. Human galectin-3 C- and N-epitope pools were immunoprecipitated from old donor sera and applied to cardiomyocytes at a 1:40 dilution. Parallel treatments involved pre-mixing these epitopes with 34 mM lactose or 100 µM gallic acid (GA) before cell incubation. **(A, B)** C-epitope formed distinct surface clusters, which were absent when pre-incubated with lactose or GA, while N-epitope exhibited a diffuse surface distribution. **(C, D)** C-epitope induced increased F-actin stress fibers and cell surface area (hypertrophy), effects blocked by lactose or GA. **(E–G)** Upregulation of RhoA-ROCK1 signaling was observed with C-epitope treatment but attenuated by lactose or GA. **(H, I)** Loss of membrane tension buffering structures, caveolae, was visualized via reduced PLA signal intensity between CAV-1 and PTRF, as PTRF dissociated from CAV1 during caveolae disassembly. **(J, K)** Increased nuclear localization of YAP1, a mechanotransduction transcription factor, was induced by C-epitope and mitigated by lactose or GA. **(L, M)** Nuclear membrane stiffening was evidenced by increased Lamin-A levels. **(N, O)** Elevated DNA damage was shown by intense γ phospho-H2AX staining. **(P–S)** Expression of hypertrophy-associated genes NFATC2 and MEF2 was increased. **(T, U)** Cardiomyocyte permeability increased, as measured by extracellular levels of LDHA (an endogenous enzyme), in culture media. Numerical keys on the X-axis: 1 =Gal-3 C- or N-epitope enriched pool, whichever applicable (generated through IP from old human donor), 2 = Gal-3 C- or N-epitope enriched pool premixed with lactose (+ Lactose/+L), 3 = Gal-3 C- or N-epitope enriched pool premixed with gallic acid (+ Gallic acid/+GA). Statistical significance was assessed by t-tests: *p ≤ 0.05, **p ≤ 0.01, ***p ≤ 0.001.

Taken together, these findings underscore the essential role of the galectin-3 C-epitope CRD in mediating cardiomyocyte hypertrophy and suggest that its inhibition by small molecules such as GA may serve as a promising therapeutic approach. A graphical summary of these results demonstrates the role of excessive galectin-3 C-epitope oligomers as critical drug targets in cardiac hypertrophy **(Figure 12)**.

**Figure 12.**
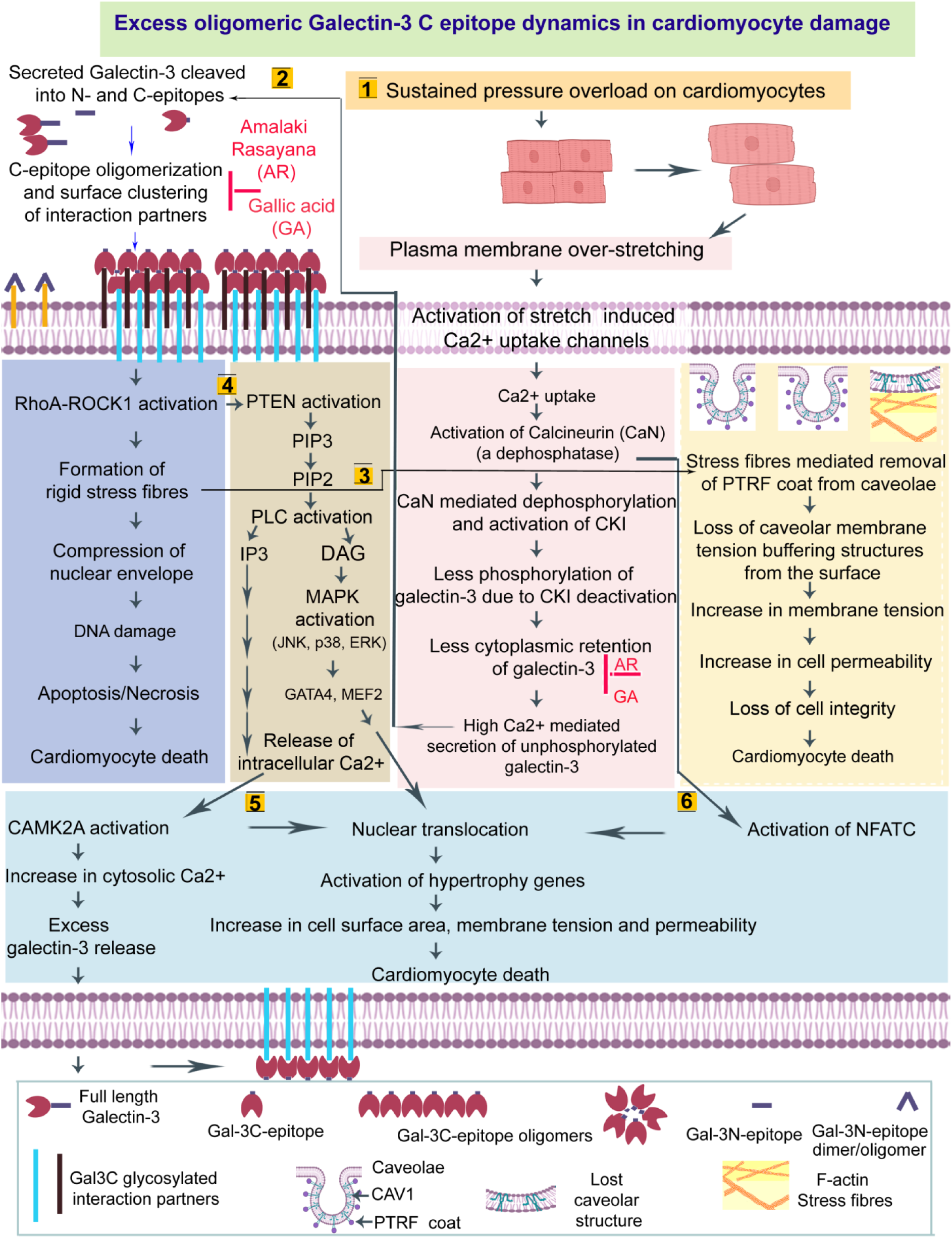
Schematic illustration of the putative mechanisms by which elevated levels of extracellularly oligomerized galectin-3 C-epitopes contribute to cardiac hypertrophy (CH) and represent key therapeutic target. 1. During CH, cardiomyocyte plasma membranes undergo dynamic hydrostatic pressure-induced stretching. This mechanical stretch activates calcium-conducting pressure/stretch sensors, increasing intracellular calcium influx. Elevated calcium activates calcineurin A (CaNA), a phosphatase that dephosphorylates casein kinase 1 (CKI). CKI normally phosphorylates galectin-3 at serine residues (notably serine 6 in the N-domain), a modification that promotes intracellular retention of galectin-3 and limits its secretion. However, CaNA-mediated CKI inactivation reduces galectin-3 phosphorylation, leading to increased secretion of unphosphorylated galectin-3 into the extracellular space. **2.** Extracellular galectin-3 is proteolytically cleaved into N- and C-domains by enzymes such as matrix metalloproteinases. These domains homo-oligomerize and multimerize on the cardiomyocyte surface, with C-epitope oligomers forming large clusters that induce intense biomechanical stress. This mechanical stress activates the RhoA-ROCK1 pathway, promoting the formation of rigid F-actin stress fibers that help withstand membrane tension but compress and deform the nucleus, causing chromatin damage. **3.** These stress fibers also cause the disassembly of caveolae—the cell’s primary membrane tension-buffering structures—by removing their PTRF coat, increasing membrane tension, loss of permeability, and ultimately cell death. **4.** The RhoA-ROCK1 pathway further activates PTEN phosphatase, converting PIP3 to PIP2. PIP2 stimulates PLC, triggering DAG-MAPK signaling and nuclear translocation of hypertrophy-associated transcription factors (MEF2, GATA4). PLC also promotes IP3-mediated calcium release from intracellular stores, amplifying intracellular calcium and calcineurin A activation, which drives further secretion of unphosphorylated galectin-3. **5.** Elevated calcium also activates CAMK2A autophosphorylation, causing additional calcium release from intracellular stores and CAMK2A nuclear translocation, which promotes hypertrophy gene expression. **6.** High calcium levels, through calcineurin A, dephosphorylate NFATC2, facilitating its nuclear translocation and upregulation of hypertrophy genes, resulting in cardiomyocyte overstretch and death. **Therapeutic mechanism:** AR extract and its bioactive component gallic acid (GA) bind to the carbohydrate recognition domain (CRD) of galectin-3, inducing a conformational change that enhances CKI accessibility to the galectin-3 N-domain. This increases galectin-3 phosphorylation, promoting intracellular retention and reducing harmful extracellular secretion. Additionally, AR and GA bind extracellular galectin-3 CRD, blocking its surface clustering and the subsequent activation of the RhoA-ROCK1-PTEN-PLC-Ca²⁺ signaling cascade, thereby mitigating calcium dysregulation and cellular damage. This figure synthesizes knowledge from the literature (citations 33, 34, 40–44) together with the results presented in this study.

### Galectin-3 C-epitope, ANP, PPIA, and ALB are promising proteins for a biomarker panel aimed at evaluating drug efficacy

Recognizing that galectin-3 is co-secreted with several proteins, we investigated their potential inclusion in a multiplexed biomarker panel, alongside the galectin-3 C-epitope, to improve the clinical assessment of drug response. Atrial natriuretic peptide (ANP) [45], peptidyl-prolyl isomerases A (PPIA)[46, 47], and albumin (ALB)[48, 49] are known to be secreted into the serum either freely or encapsulated in extracellular vesicles such as exosomes and microvesicles, in left ventricular hypertrophy.

Therefore, ANP, PPIA, and ALB were prioritized based on their functional association with cardiac pathology. We propose combining galectin-3 detection with natriuretic peptides like ANP—secreted in abundance by mechanically stressed cardiomyocytes—to enhance correlation between serum galectin-3 levels and cardiac tissue pathology **(Figure 9)**. Additionally, PPIA and ALB are implicated in cardiovascular dysfunction and were found to be dynamically regulated in our models.

In the pressure overload-induced cardiac hypertrophy (PO-CH) rat model, animals treated with Amalaki rasayana (AR) for at least 12 months showed a significant reduction in serum levels of ANP, PPIA, ALB, and the galectin-3 C-epitope compared to untreated animals **(Figure 13).** Similarly, aged (21-month-old) Wistar rats treated with AR also displayed lower levels of these proteins than age-matched untreated controls, supporting the preventive therapeutic efficacy of AR.

**Figure 13:**
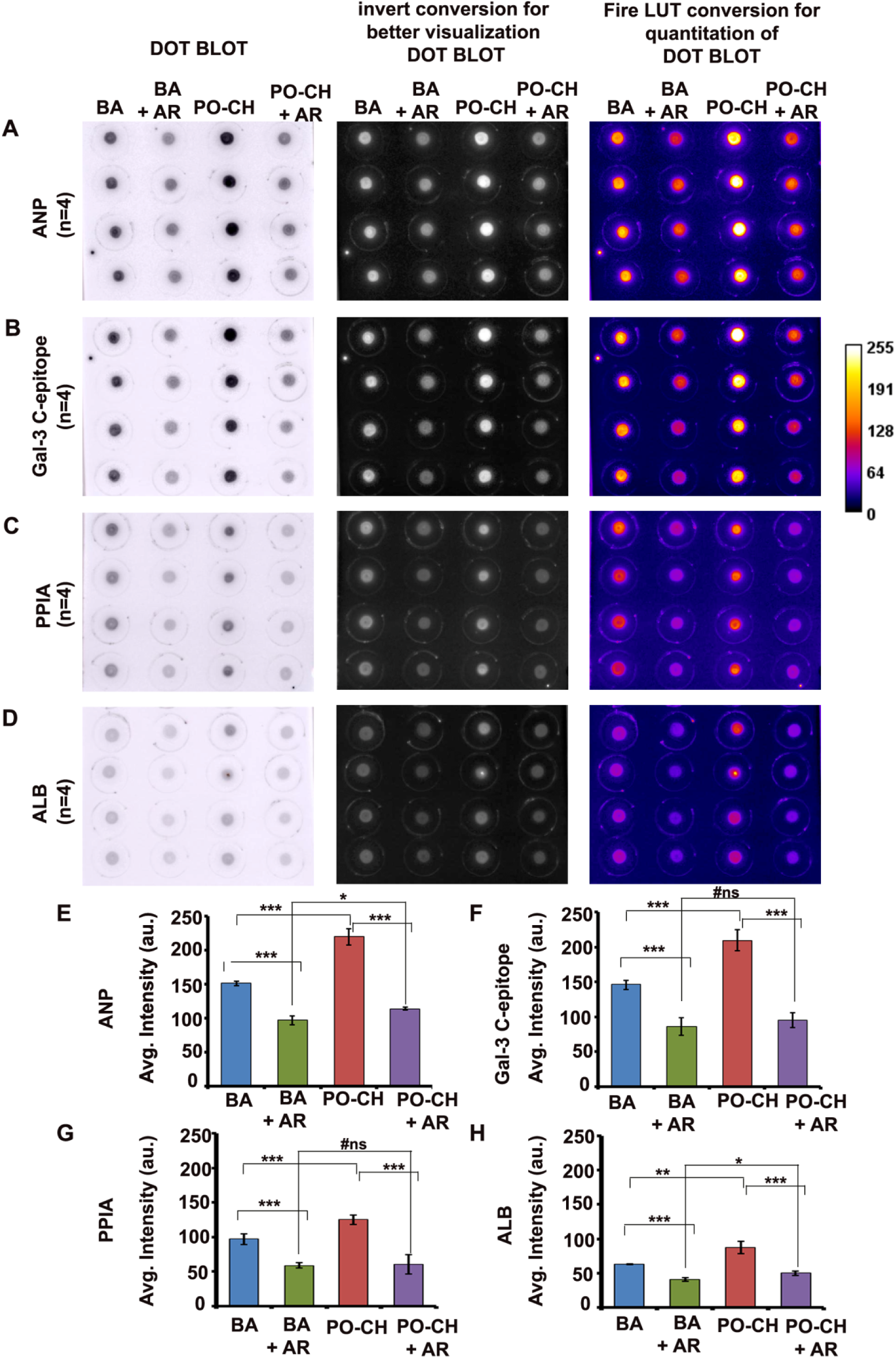
Correlated reduction of galectin-3 C-epitope levels with ANP, PPIA, and ALB in AR-treated rats. Dot blot assays measuring **(B, F)** galectin-3 C-epitope and its co-secretory partners—**(A, E)** atrial natriuretic peptide (ANP), **(C, G)** peptidylprolyl isomerase A (PPIA), and **(D, H)** albumin (ALB)—in serum samples from BA, BA+AR, PO-CH, and PO-CH+AR rat cohorts (n=4 per group). Each dot corresponds to 6 µg of serum protein and was probed using knockout-validated antibodies specific to each target. Data are presented as mean ± SD, with statistical significance assessed by t-tests; *p ≤ 0.05, **p ≤ 0.01, ***p ≤ 0.001

To evaluate the translational potential of this biomarker panel in humans, we conducted Western blot analyses using serum samples from healthy donors and hypertrophic cardiomyopathy (HCM) patients on drug therapy. Consistent with rodent findings, older donors and HCM patients exhibited higher levels of ANP, PPIA, and ALB than younger individuals **(Figure 9, 14)**. Importantly, the levels of ANP, PPIA, and ALB were comparable between older healthy donors and treated HCM patients. However, C-epitope oligomer levels were modestly lower in the HCM cohort than in healthy age-matched individuals, indicating that C-epitope reduction could serve as a marker of treatment efficacy (Figure 9 and Figure 14).

**Figure 14:**
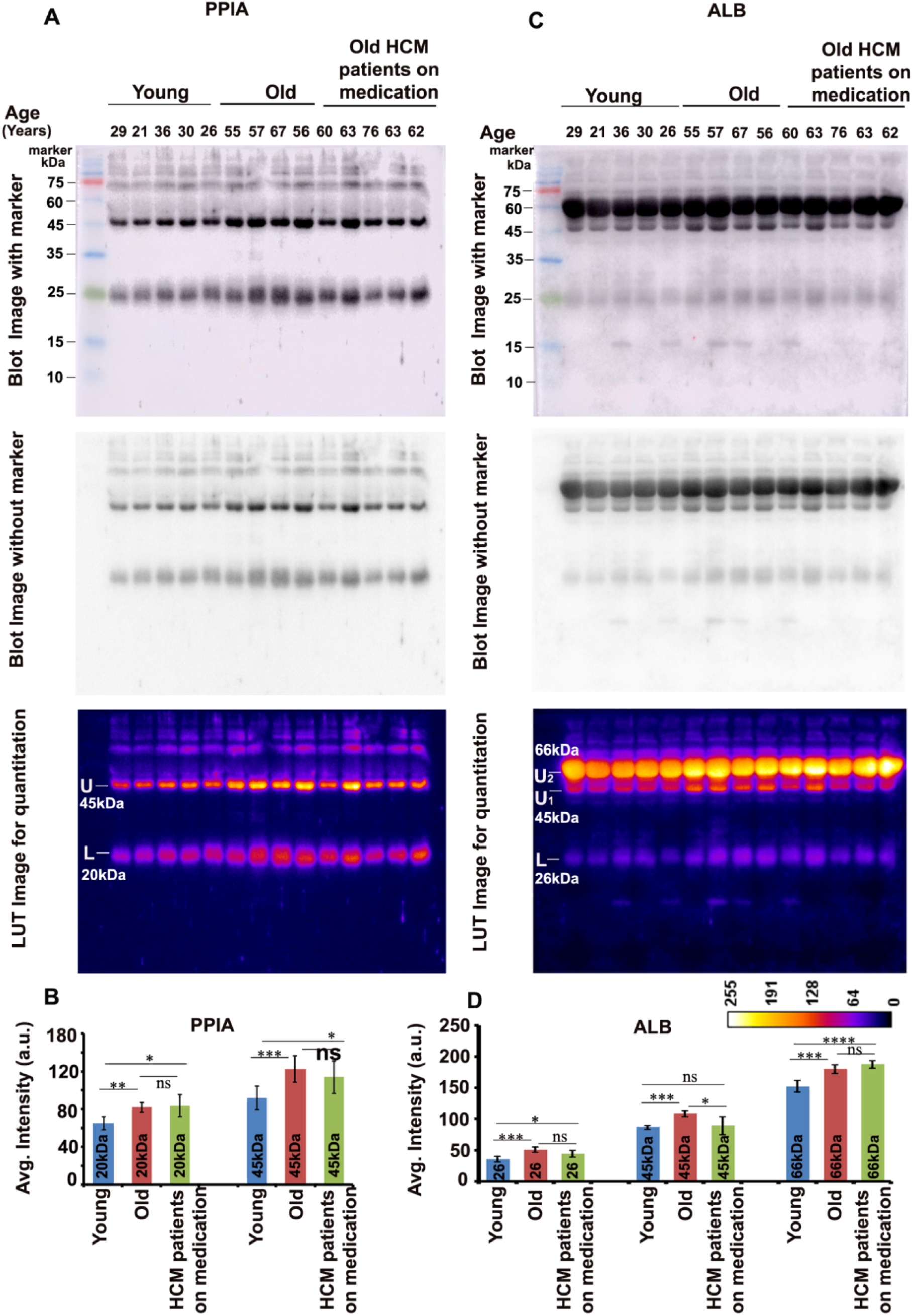
Quantification of PPIA and ALB levels in human sera. **(A-D)** Serum samples from young individuals, older adults, and cardiac hypertrophy survivors receiving diverse pharmacological treatments were analyzed for peptidylprolyl isomerase A (PPIA) and albumin (ALB) levels. Data are shown as mean ± SD, with statistical significance determined by t-tests; *p ≤ 0.05, **p ≤ 0.01, ***p ≤ 0.001

Furthermore, donor samples exhibiting high levels of C-epitope oligomers also demonstrated elevated concentrations of PPIA and ALB **(Figure 14)**, reinforcing the correlated secretory pattern of these proteins with that of galectin-3. These findings support the inclusion of galectin-3 C-epitope, ANP, PPIA, and ALB in a multiplexed serum biomarker panel for the monitoring of drug response and disease progression in CH.

Ongoing work is focused on quantifying the dynamic concentration ranges of these biomarkers across larger and regionally diverse human cohorts. This data will be instrumental in designing immunoassay and microfluidics-based biosensors for improving the predictive accuracy and clinical utility of biomarker-based monitoring of drug efficacy in heart failure.

## DISCUSSION

In the context of aging-associated and pressure overload-induced left ventricular hypertrophy (PO-CH/LVH), our findings present compelling evidence that the galectin-3 C-terminal carbohydrate recognition domain (C-epitope) oligomers represent a more sensitive and functionally relevant biomarker than full-length galectin-3 or its N-terminal counterparts.

Both in serum from aged human donors and in a pressure overload–cardiac hypertrophy (PO-CH) animal model, we observed a significant increase in circulating C-epitope oligomers. These changes correlated with elevated levels of atrial natriuretic peptide (ANP)—a known marker of myocardial stretch—and other co-secreted proteins such as cyclophilin A (PPIA) and albumin (ALB), suggesting a concerted secretory stress response associated with myocardial remodeling [45–49].

Therapeutic intervention with Amalaki rasayana (AR), an Ayurvedic nutraceutical-based medicine enriched in gallic acid, markedly reduced serum levels of galectin-3 C-epitope oligomers, ANP, PPIA, and ALB. These reductions were associated with structural and functional cardiac improvement in PO-CH models, positioning C-epitope oligomer profiling as a reliable indicator of therapeutic efficacy. Our data support the concept that monitoring galectin-3 C-epitope and its co-secreted partners can serve not only to gauge clinical recovery in heart failure patients, but also to stratify risk and monitor prophylactic responses, particularly in aging populations predisposed to cardiac stress [50, 51].

Mechanistically, AR appears to exert its cardioprotective effects in part through gallic acid’s high-affinity binding to the C-epitope CRD. This interaction competitively inhibits galectin-3’s glycan-dependent surface binding, thereby blocking downstream hypertrophic mechanotransduction via RhoA-ROCK1 signaling and stress fiber formation. Moreover, CRD binding by gallic acid facilitates conformational changes that expose galectin-3 to phosphorylation, promoting its cytoplasmic retention and reducing extracellular secretion— key steps in mitigating maladaptive cardiac remodeling [16, 33].

Importantly, serum analyses from hypertrophic cardiomyopathy (HCM) patients receiving pharmacological treatment revealed lower levels of galectin-3 C-epitope oligomers compared to age-matched healthy donors, despite having higher total galectin-3 levels. This uncoupling of total and functional galectin-3 levels underscores the inadequacy of conventional full-length galectin-3 ELISA platforms in accurately reflecting therapeutic impact. Our data show that C-epitope levels, along with PPIA, ALB, and ANP, form a synergistic network that better reflects disease progression and resolution than individual biomarkers alone.

These findings advocate for broader adoption of a multiplex biomarker panel, incorporating galectin-3 C-epitope, PPIA, ALB, and ANP, to improve clinical stratification, prognosis, and therapy response assessment in heart failure. We propose the development of low-cost, paper-based multiplex diagnostic tools, high-sensitivity ELISA assays, and reverse-phase protein array (RPPA) chips specifically designed to detect these markers in diverse patient populations. Furthermore, clinical validation of AR’s efficacy across large, geographically distributed cohorts is urgently needed to establish its safety and translational potential in cardiometabolic health.

Beyond cardiac hypertrophy, elevated levels of galectin-3 and its C-epitope oligomers have been implicated in a spectrum of diseases including malignancies, non-alcoholic steatohepatitis (NASH), chronic kidney disease, obesity, hypertension, inflammatory bowel disease, dementia, diabetes, and even COVID-19[52–53]. Our study suggests that quantifying galectin-3 C-epitope enrichment may offer a sensitive, functionally relevant window into systemic disease states and therapeutic trajectories, offering promise for precision medicine approaches in both cardiovascular and multisystem disorders.

## CONCLUSIONS

Galectin-3 performs multiple vital physiological functions, and thus, complete inhibition or dysregulation of its endogenous and exogenous levels risks significant adverse effects [13, 16]. Our research distinctly highlights the pivotal role of galectin-3 C-epitope oligomeric forms as superior, highly specific biomarker for the diagnosis and therapeutic monitoring of hypertrophic cardiomyopathy, surpassing the diagnostic value of full-length galectin-3 protein.

These findings underscore the urgent need for the development of accessible, low-cost, paper-based biomarker assays incorporating galectin-3 C-epitope in combination with PPIA, ALB, and ANP, enabling sensitive and reliable clinical evaluation of cardiac hypertrophy and drug efficacy.

Moreover, our work identifies the excessively secreted galectin-3 C-epitope oligomers as a promising and actionable drug target in age-associated and pathological cardiac hypertrophy, with Amalaki rasayana demonstrating significant therapeutic potential to suppress these harmful oligomeric pools and ameliorate cardiac remodeling **(Figure 15)**.

**Figure 15:**
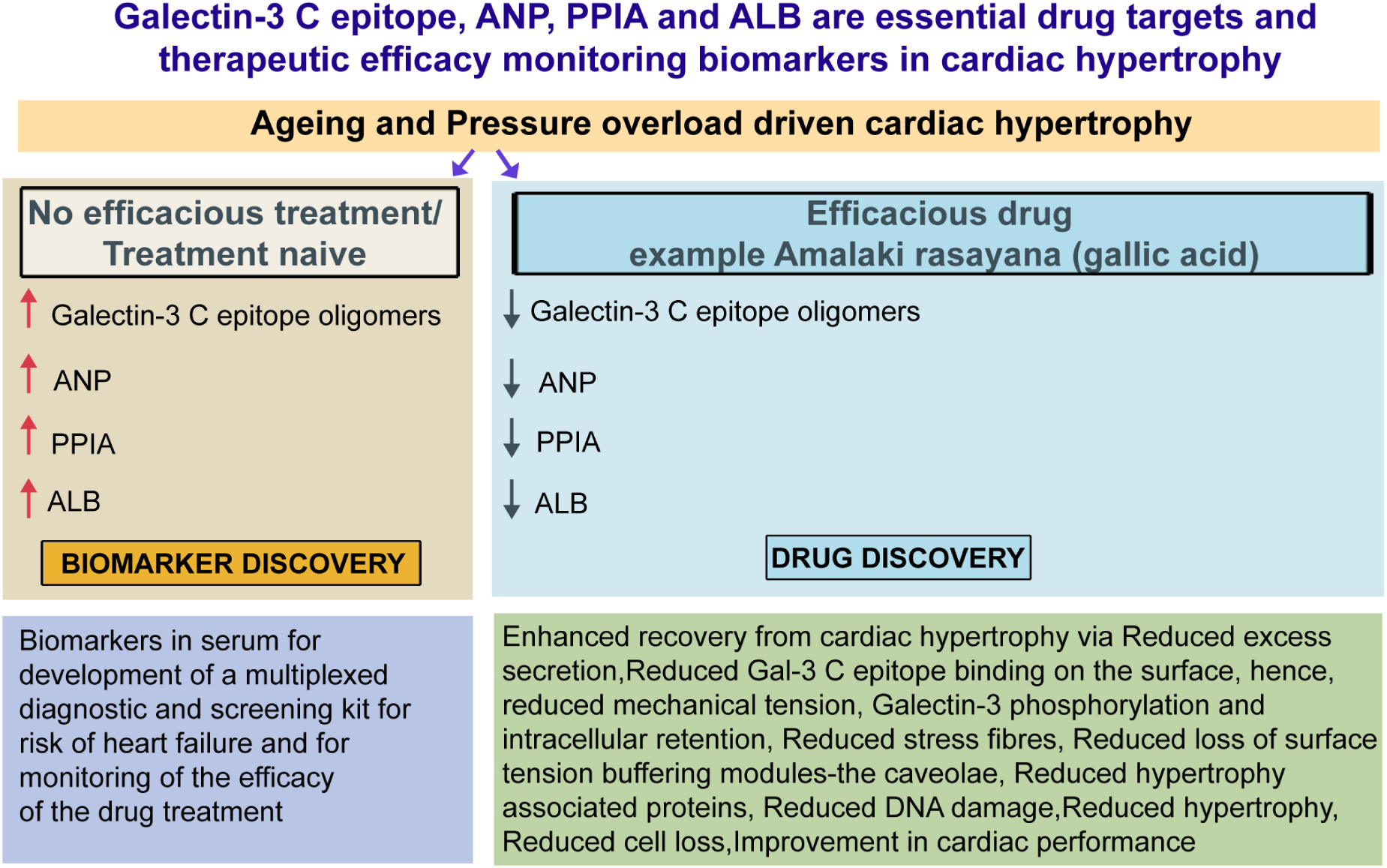
Summary of the research findings. Aging and pressure overload-driven cardiac hypertrophy result in elevated tissue and circulating levels of galectin-3 C-epitope oligomers, ANP, PPIA, and ALB—key proteins implicated in cardiac damage. Treatment with Amalaki rasayana (AR) or gallic acid (GA) leads to significant improvements in cardiac function, morphology, cellular and molecular parameters, alongside marked reductions in galectin-3 C-epitope oligomers, PPIA, ANP, and ALB levels. These proteins collectively serve as a biomarker panel to monitor therapeutic efficacy. Additionally, AR/GA-mediated inhibition of excessive extracellular galectin-3 C-epitope oligomers attenuates cellular tension, RhoA-ROCK1 signaling, formation of rigid F-actin stress fibers, loss of caveolae, hypertrophy gene expression, cardiomyocyte surface area, DNA damage, and cell death.

Collectively, these insights pave the way for geroprotective and precision cardiovascular medicine approaches focused on refined biomarker panels and targeted therapeutics to mitigate cardiac hypertrophy and improve patient outcomes.

## MATERIALS AND METHODS

### Reagents, Cell Lines, and Patient Samples

Cell lines H9c2(2-1), HEK-293 (293), and CRL-1446 were procured from ATCC, Sigma, or Lonza, unless otherwise stated. The pEGFP-hGal3 plasmid (#73080) was obtained from Addgene (USA); other constructs were custom-synthesized by Zellebiotech.com. Most reagents were purchased from Merck/Sigma-Aldrich. Galectin-3 ELISA kits—Rat ERLGALS3 and Human DGAL30—were sourced from Thermo Scientific and R&D Systems, respectively. Antibodies and dilutions used are as follows:CBP-112 AP, Rabbit anti-Galectin-3 (C-epitope): 1:1000;CBP-101 AP, Rabbit anti-Galectin-3 (N-epitope): 1:1000;Galectin-3 Mouse mAb (B2C10, sc-32790): 1:500; A13506, Galectin-3 KO validated ab 1:1000; NPPA Rabbit pAb (A1609): 1:5000; Albumin Rabbit pAb (A1363): 1:1000;Anti-Cyclophilin A (PPIA, AB58144): 1:5000. Human heart tissue arrays were acquired from TissueArray.Com (USA). Human serum samples were sourced from Innovative Research (USA) following approval from the Institutional Human Ethics Committee of RGCB (File no. IHEC/12/2023_Ex/07) and in accordance with ICMR guidelines, Government of India.

### Development of a Drug Model for Cardiac Hypertrophy

To validate a biomarker panel for assessing therapeutic efficacy in hypertrophic cardiomyopathy (CH), a preclinical drug model with established anti-hypertrophic effects was employed [18]. All animal procedures were approved by the Institutional Animal Ethics Committee (IAEC) of RGCB and conducted in accordance with guidelines of the Committee for Control and Supervision of Experiments on Animals (CCSEA), Government of India.

Three-month-old male Wistar rats (180–200 g) were housed under standard conditions (12 h light/dark cycle, controlled temperature and humidity) with ad libitum access to synthetic chow and water. Cardiac hypertrophy was induced by transverse aortic constriction (TAC) using a titanium clip to reduce aortic diameter by ∼60%. Anesthesia was administered using 3% isoflurane in 100% oxygen. Aortic constriction was verified via trans-thoracic 2D colour Doppler imaging. Sham-operated controls (BA and BA+AR) underwent identical surgery without clip placement. Left ventricular hypertrophy was monitored from nine months of age.

*Two experimental arms were included:*

1. *Aging Group:* Sixteen rats were divided into ‘BA’ (biologically aged, sham control) and ‘BA+AR’ groups. The BA+AR group received Amalaki rasayana (AR) orally (500 mg/kg, 5 days/week) starting at 9 months, continued until 21 months. Dose selection was based on prior efficacy and traditional use.
2. *LVH Group:* Sixteen rats underwent TAC at 3 months. From 9 months, one cohort received AR (‘PO-CH+AR’), while the other remained untreated (‘PO-CH’). Treatment continued until 21 months of age.

This model effectively simulates age-related and pressure overload-induced cardiac hypertrophy for evaluating therapeutic interventions.

### Drug Administration

Amalaki rasayana (AR) was prepared and characterized as previously described [18]. It was administered orally each morning, 5 days per week, with a 2-day interval to allow systemic clearance of excess components. In both the aging and pressure overload-induced cardiac hypertrophy (PO-CH) groups, treatment continued until 21 months of age, ensuring at least 12 months of therapeutic exposure.

### Echocardiography

Echocardiographic evaluations were conducted at multiple time points before and during treatment, up to 21 months of age, in both BA and PO-CH groups. Left ventricular dimensions and ejection fraction were measured to assess the impact of AR on cardiac changes associated with aging and pressure overload-induced hypertrophy.

### Exercise Tolerance

One month before study completion, all rats underwent a standardized treadmill protocol (30 min/day, 5 days/week). Tolerance was assessed by recording fatigue time and distance covered at incremental speeds of 5, 10, and 15 m/min.

### Tissue Collection

At the end of the study, rats were euthanized, and cardiac tissues were collected. Samples were either fixed in 10% neutral-buffered formalin for histology or stored in RNAlater for downstream analyses, including Western blotting and qPCR. Formalin-fixed tissues were processed for paraffin embedding.

### Assessment of Cardiac Fibrosis, Calcification and Hypertrophy

Cardiac fibrosis was assessed in 5 µm heart sections stained with Picrosirius Red (#365548, Sigma). Slides were incubated in stain for 1 h, rinsed with distilled and acidified water, dehydrated in graded ethanol and xylene, and mounted with DPX. Fibrotic area was quantified using Fiji software and expressed as a percentage of total tissue area. Cardiomyocyte hypertrophy was evaluated by measuring myocyte dimensions in Fiji software.

Calcification was assessed using 1% Alizarin S (pH 4.2) staining. Sections were incubated for 30 min at room temperature, rinsed, and then briefly immersed in acetone and an acetone– xylene (1:1) mixture before clearing in xylene and mounting with DPX. Calcium deposits appeared red to orange and were quantified using image analysis software.

### Immunohistochemistry

Slides were pre-baked at 55 °C for 15 min, de-paraffinized in xylene (2 × 20 min), and treated with chloroform. Rehydration was performed through a graded ethanol series (100% to 70%, 5 min each). Sections were washed in PBS (pH 7.4) and permeabilized with 0.01% digitonin in PBS for 30 min at room temperature.

Blocking was done using 3% BSA and 2% donkey serum in PBS for 1.5 h, followed by overnight incubation with primary antibodies at 4 °C. After PBS washes, secondary antibodies were applied for 1 h at room temperature. Nuclei were counterstained with DAPI (5 min), and slides were mounted with 70% glycerol. Imaging and analysis followed previously established protocols.

### Cell Culture and Conditions

Cardiomyoblasts were cultured in low-glucose DMEM supplemented with 10% FBS and antibiotics (complete medium). For experiments, cells were seeded in 6-well plates, 8-well chamber slides, or 96-well plates at densities of 0.1, 0.01, or 0.001 million cells/well, respectively. After 24 h, differentiation into myocytes was induced by switching to serum-reduced medium (DMEM with 1% FBS and antibiotics) for another 24 h. Cells were then treated in complete medium containing 5% FBS.

### MTT Assay for Dose Estimation of AR

Cardiomyoblasts (5,000 cells/well) were seeded and differentiated in 96-well plates. Post-differentiation, cells were treated with varying concentrations of AR (0, 5, 10, 50, 150, 200 µg/mL) in complete medium for 24–48 h. After treatment, cells were washed with warm 1× HBSS, and MTT (0.5 mg/mL) was added to each well. Plates were incubated in the dark at 37 °C for 4 h. Formazan crystals were solubilized with 100 µL DMSO/well and gently rocked for 15 min. Absorbance was recorded at 570 nm using a Varioskan multimode plate reader (Thermo Scientific, USA). Cell viability was expressed as the OD (optical density) in treated versus untreated cells.

### Pressure Overload Cellular Model

A pressure overload model was established using a weight-based method [26, 27]. Cardiomyoblasts were cultured in low-glucose DMEM with 10% FBS and antibiotics in 6-well plates. After 24 h, differentiation was induced by incubating cells in 1% FBS medium for 24 h. At ∼80% confluence, medium was replaced with CO₂-independent medium containing 5% FBS to minimize pH fluctuations under reduced gas exchange.

Mechanical stress was simulated by placing sterile 10 g weights (∼1 g/cm²) on the medium surface, applying consistent pressure to mimic pathological cardiomyocyte overload. Cells were exposed to pressure for 12–48 h; controls (NO-PO) were cultured without weights but covered with dummy lids to ensure equal gas exchange. PO enhanced the cell surface area of cardiomyocytes by 3-4 fold, akin to that observed in the in vivo animal experiments.

Following treatment, three independent experiments per group (NO-PO and PO) received AR (100 µg/mL) or GA (100 µM), as per doses determined via viability assays; while three experimental sets remained untreated. Cells were harvested for analysis of hypertrophic markers, protein expression, signaling pathways, and validation of mechanical load–induced hypertrophy.

### Dot Blot ELISA

Proteins were extracted using lysis buffer and quantified by BCA assay to normalize concentrations. Equal amounts (6 µg in 2.2 µ L) were spotted onto 0.45 µm nitrocellulose membranes (Amersham), air-dried for 2 h at room temperature, and blocked overnight at 4 °C with 3% BSA in TBST (0.1% Tween-20). Membranes were washed thrice with TBST, incubated with primary antibodies diluted in blocking buffer for 1 h, followed by HRP-conjugated secondary antibodies for 1 h at room temperature. Signals were detected using enhanced chemiluminescence and imaged digitally. Dot intensities were quantified with Fiji software to assess protein expression.

### Human Galectin-3 ELISA

Serum samples were diluted 1:1 and processed as per the manufacturer’s protocol. Absorbance was measured at 450 nm using a Varioskan multimode plate reader (Thermo Scientific, USA). Data represent mean ± SD from three independent experiments.

### Immunocytochemistry

Cells were fixed in 1.5% paraformaldehyde for 20 min, permeabilized with 0.25% saponin for 20 min, and blocked for 1 h in PBS containing 3% BSA and 2% normal donkey serum. Primary antibodies were applied overnight at 4 °C. After PBS washes, cells were incubated with Alexa Fluor–conjugated secondary antibodies (1:200; Jackson ImmunoResearch) for 1 h at room temperature in the dark. Slides were mounted with DAPI-containing medium (Invitrogen), and images were captured using an Olympus confocal microscope (60× oil objective, NA 1.29). Imaging and analysis followed established protocols. Duolink in situ proximity ligation assay was performed according to the manufacturer’s instructions (Sigma, USA)[54].

### Western Blotting

Cells were washed with 1× PBS and lysed in RIPA buffer (0.1% NP-40, 0.1% SDS, 0.5% sodium deoxycholate, 150 mM NaCl, 50 mM Tris-HCl pH 7.4, protease inhibitors). Lysates were sonicated and centrifuged at 14,000 rpm for 15 min at 4 °C; supernatants were collected. Equal amounts of protein were mixed with SDS-β-mercaptoethanol sample buffer, heated at 95 °C for 5 min, separated by SDS-PAGE, and transferred to PVDF membranes.

Membranes were blocked overnight in TBST (0.1% Tween-20) with 3% BSA, then incubated with primary antibodies diluted in blocking buffer for 1 h at room temperature or overnight at 4 °C. After washes, membranes were probed with HRP-conjugated secondary antibodies and developed using Clarity ECL (Bio-Rad). Signals were detected with LAS-500 imaging system.

### Plasmid Transfection

Cells at 70% confluence were incubated in serum-free OPTIMEM media for 2 h prior to transfection. High-purity plasmids (0.5 µg) were complexed with Lipofectamine LTX Plus reagent at room temperature for 15–30 min. The mixture was then diluted with 350 µ L OPTIMEM and added dropwise to cells with gentle swirling. After 4–6 h incubation at 37 °C, 5% CO₂, the transfection medium was replaced with complete growth medium. Cells were cultured for 24–72 h for optimal expression.

### Galectin-3 WT and Mutant Conditioned Medium Experiments

HEK293 cells were transfected with Gal-3 EGFP wild-type or carbohydrate recognition domain mutant Gal-3 R186S EGFP plasmids. Following pressure overload treatment, conditioned media containing secreted Gal-3 were collected. Cardiomyocytes were then incubated with either Gal-3 EGFP or R186S EGFP media for 24 h. Cells were fixed in 1.5% PFA for surface imaging or stained with rhodamine phalloidin to visualize F-actin stress fibers and changes in the cardiomyocytes’ surface area.

### Rhodamine Phalloidin and WGA Staining

Fixed cells, prepared for immunocytochemistry if needed, were incubated with Rhodamine phalloidin (2 µg/mL) for 45 min to visualize F-actin stress fibers. For WGA staining, cells were treated with WGA (5 µg/mL) for 45 min, followed by brief washes. Cells were counterstained with Hoechst (5 µg/mL, 5 min) and mounted. Imaging parameters and analysis were consistent with prior protocols [54].

### Cell Surface-Bound Galectin-3 Extraction via Acid Wash

At the endpoint of PO and AR/GA treatments, media were collected, and plates with live cells were placed on ice. Cells were incubated with complete medium adjusted to pH 2.0 for 45 minutes to strip surface-bound galectin-3. Released gal-3 was analyzed from the acid wash media normalized to normal pH, while cells were lysed separately to assess intracellular gal-3 levels.

### Human Serum Conditioned Media Experiment

Human serum was diluted in cardiomyocyte culture medium at 1:20, 1:40, and 1:60 ratios. Differentiated cardiomyocytes were incubated with these dilutions for 10 hours before fixation. Samples were then processed to assess cell surface area and detect surface-bound N- and C-epitope galectin-3 by antibody staining.

### Immunoprecipitation and Epitope Enrichment from Human Serum

Selective enrichment of N- or C-epitope galectin-3 from aged human serum samples (n=3) was performed using Dynabeads Protein A immunoprecipitation according to the manufacturer’s protocol. Dynabeads (1.5 mg) were conjugated with 4 µg of anti-N- or anti-C-epitope galectin-3 antibodies. A total of 1600 µg serum protein (20 µl serum diluted in 580 µl PBST) was incubated overnight at 4°C with antibody-conjugated beads on a rotator. The following day, supernatants were collected, and 225 µl aliquots were incubated overnight at 4°C with or without 34 mM lactose.

Lactose-treated and untreated supernatants (225 µl each) were mixed 1:1 with cardiomyocyte culture medium, and cardiomyocytes were incubated for 10 hours. Cells were then fixed, and surface area was analyzed by capturing five random images per condition, quantifying approximately 200 cells.

Equal volumes of supernatants were also analyzed by Western blot to confirm N- and C-epitope enrichment, with the expectation of reciprocal enrichment (e.g., N-epitope bead pull-down enriched for C-epitope in the supernatant).

### Surface Visualization of N- and C-Epitope of Galectin-3

To visualize surface-bound N- and C-epitope pools of galectin-3, fixed cells were incubated with the respective antibodies for 45 minutes without permeabilization. For simultaneous visualization of both epitopes on the same tissue section, primary antibodies (N-epitope-FITC conjugated, C-epitope Alexa 594 conjugated) were applied sequentially, and signals were developed.

### Galectin-3 Docking and Simulation Studies with AR Components

The carbohydrate-binding site of human Galectin-3 (Gal-3, PDB ID: 3ZSK) and a homology-modeled structure of rat Gal-3 were prepared using the Schrödinger Protein Preparation Wizard (Schrödinger Release 2021-3: Glide, Schrödinger, LLC, New York, NY, 2021). The aim was to determine the binding conformations and energies of 18 compounds identified in the Amalaki rasayana (AR) extract. Receptor structures were energy-minimized with the OPLS_2005 force field, constraining heavy atoms to deviate no more than 0.3 Å from their original positions.

For molecular docking, Glide [55, 56, 57] was used to generate receptor grids centered on the ligand-binding site defined by the human Gal-3 crystal structure, ensuring accurate representation of the binding pocket. These grids served as docking targets for the selected compounds.

The LigPrep module generated 3D structures of the 18 compounds, optimized their geometry, and produced all possible stereoisomers and ionization states. The energy-minimized ligands were docked into the receptor grids using Glide in standard precision (SP) mode with pre-defined constraints.

The docking protocol predicted the preferred conformation and orientation of each ligand within the receptor’s binding site. The pose with the lowest GlideScore (GScore) was selected as the best-docked conformation and used for subsequent interaction analysis.

### Statistics

All experiments were performed in triplicate, with error bars representing the standard deviation (SD) calculated from three independent experiments. Statistical significance for pairwise comparisons was determined using Bonferroni’s t-test, with significance levels indicated as *p ≤ 0.05, **p ≤ 0.01, and ***p ≤ 0.001. Data are presented as means ± SD from a minimum of three independent trials.

For single-cell analyses, between 50 and 200 cells were quantified from five randomly selected fields per condition, across three independent experiments. Image analysis was conducted using Fiji software, and calibration bars for LUT-converted images are included in the corresponding figures.

## DECLARATIONS

### Ethics approval and consent to participate

Not applicable.

### Consent for publication

Not applicable.

### Availability of data and materials

All data generated during the current study are available in the manuscript itself, including un-cropped, original western and dot representative images.

### CRediT authorship contribution statement

**Puja Laxmanrao Shinde:** Methodology, Formal analysis, Investigation, Visualization. **Vikas Kumar:** Methodology, Formal analysis. **Siddhartha Singh:** Methodology, Formal analysis, Investigation, Visualization. **K.C. Sivakumar :** Methodology, Formal analysis, Investigation. **Rashmi Mishra:** Conceptualization, Methodology, Formal analysis, Resources, Investigation, Visualization, Supervision, Project administration, Funding acquisition, Writing – review & editing.

## Acknowledgments

PLS and SS acknowledge research fellowships from the Department of Biotechnology (DBT) and University Grant Commission (UGC), respectively.

